# Transcriptome innovations in primates revealed by single-molecule long-read sequencing

**DOI:** 10.1101/2021.11.10.468034

**Authors:** Luis Ferrández-Peral, Xiaoyu Zhan, Marina Álvarez-Estapé, Cristina Chiva, Paula Esteller-Cucala, Raquel García-Pérez, Eva Julià, Esther Lizano, Òscar Fornas, Eduard Sabidó, Qiye Li, Tomàs Marquès-Bonet, David Juan, Guojie Zhang

## Abstract

Transcriptomic diversity greatly contributes to the fundamentals of disease, lineage-specific biology, and environmental adaptation. However, much of the actual isoform repertoire contributing to shaping primate evolution remains unknown. Here, we combined deep long- and short-read sequencing complemented with mass spectrometry proteomics in a panel of lymphoblastoid cell lines (LCLs) from human, three other great apes, and rhesus macaque, producing the largest full-length isoform catalog in primates to date. Our transcriptomes reveal thousands of novel transcripts, some of them under active translation, expanding and completing the repertoire of primate gene models. Our comparative analyses unveil hundreds of transcriptomic innovations and isoform usage changes related to immune function and immunological disorders. The confluence of these innovations with signals of positive selection and their limited impact in the proteome points to changes in alternative splicing in genes involved in immune response as an important target of recent regulatory divergence in primates.

## Introduction

The vast complexity of eukaryotic transcriptomes arises from the co-occurrence of multiple RNA processing events leading to the generation of different isoforms encoded by the same gene. Particularly, the fine-tuning of alternative splicing (AS) is a critical mechanism underlying disease and phenotypic evolution ^1,2^. Hence, previous research on the conservation of RNA processing events in human and non-human primates (NHP) has revealed the functional importance of transcriptomic diversity ^3–10^. Interestingly, the potential of increasingly complex transcriptomes to produce alternative protein isoforms is under intense debate as most of the predicted proteins remain undetected ^11–17^ , and alternative transcripts can carry out other regulatory functions ^18^.

Most previous comparative studies are based on short-read RNA-seq, which cannot capture the actual isoform landscapes in different species. Notably, the recent emergence of full-length isoform sequencing overcomes this limitation and has contributed to the refinement of isoform repertoires in model and non-model organisms ^19^. Thus, single-molecule sequencing in combination with short-read RNA-seq and proteomic data appears as a powerful tool to disentangle the recent primate evolution of transcriptomes with isoform resolution, especially with the most recent high-quality genome assemblies for NHP ^5,20–22^ allowing the accurate definition of orthologous regions in primates.

In this study, we inspect the recent evolutionary dynamics of splicing programs using high-quality full-length transcriptomes obtained by integrating multi-species deep Iso-seq and RNA-seq data for a panel of lymphoblastoid cell lines (LCLs) from human and NHP (Fig. 1). These full-length transcriptomes substantially expand the current isoform repertoires in primates, and additional proteomic experiments (MS/MS) provide evidence of protein translation from novel transcripts (Fig. 1). We leveraged our transcriptomes to reconstruct transcript expression gains and losses in the primate lineage, linking these expression differences to genetic changes in splice sites and novel exonization events. Innate immune system-related genes and cell proliferation genes are preferential targets of evolutionary innovations. Moreover, the convergent accumulation of these innovations and their occurrence in genes under positive selection in primates highlight the strength of the evolutionary pressures suffered by the immune response and other cell type-specific functions.

**Figure 1.**
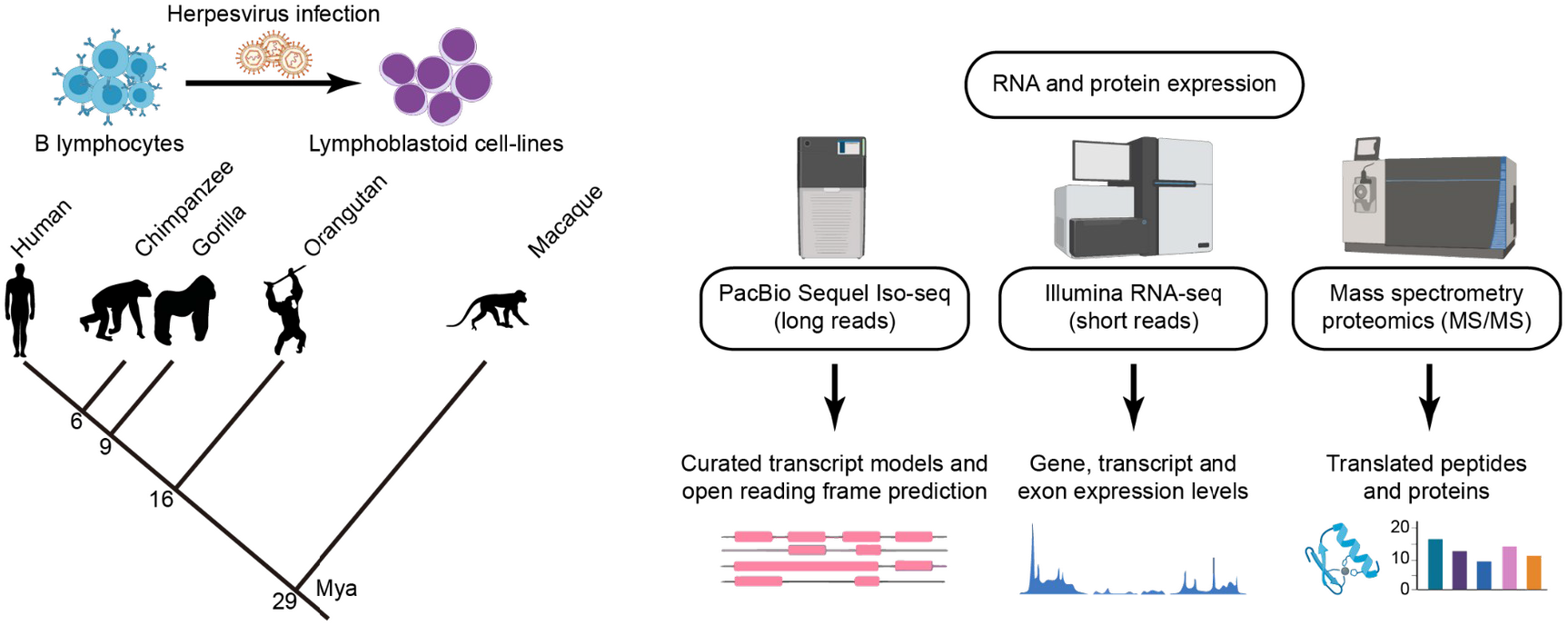
Characterization of transcriptomes and proteomes of human, chimpanzee, gorilla, orangutan and rhesus macaque lymphoblastoid cell lines (LCLs). We performed multi-species full-length isoform sequencing (PacBio Iso-seq), RNA-seq (Illumina), and tandem mass spectrometry experiments (MS/MS). Mya: million years ago.

## Results

### Isoform sequencing largely improves human and other primates annotations

To define high-quality transcriptomes, we combined high-depth long reads with short reads in our panel of LCLs derived from human and NHP, resulting in the largest full-length isoform catalog in primates to date. Briefly, Iso-seq subreads for 2 human, 2 chimpanzee, 2 gorilla, 1 (2 isogenic cultures) orangutan, and 2 rhesus macaque LCLs were processed to produce an average of over 2.3 million circular consensus sequences (CCS) per species (range from 1.9 to 2.9 million CCS). A strict quality control -including base correction and splice junction adjustment with RNA-seq- was conducted to refine the transcript models (Methods). As a result, we defined a total of 148,610 isoforms (on average, 29,722 per species) in 42,814 genes (on average, 8,563 genes per species), which were further assessed by SQANTI ^23^ (for the summary statistics of each processing step see Supplementary Table 1).

Our refined transcriptomes reveal a considerable fraction of novel isoforms (49%) derived from unreported combinations of annotated splice sites, novel splice sites, fusion events, and intergenic transcription (Fig. 2a, Supplementary Fig. 1, Supplementary Table 2 and Supplementary Data 1). Remarkably, we also expanded the transcript models recently defined by isoform sequencing for the Universal Human Reference RNA (UHRR) (Supplementary Fig. 2). The alternative splicing events newly disclosed in this study (73% of all detected events) unveil an unexpected level of isoform diversity in primates (Supplementary Fig. 3). These events are mostly associated with canonical splice sites (GT-AG, GC-AG, and AT-AC), and thus they agree with the previous knowledge on RNA splicing mechanisms. Moreover, the contribution of different AS modes is similar across all primate species, and in line with previous observations, exon skipping (SE) and retained introns (RI) are the most prevalent mechanisms ^3^ (Fig. 2b and Supplementary Table 3).

**Figure 2.**
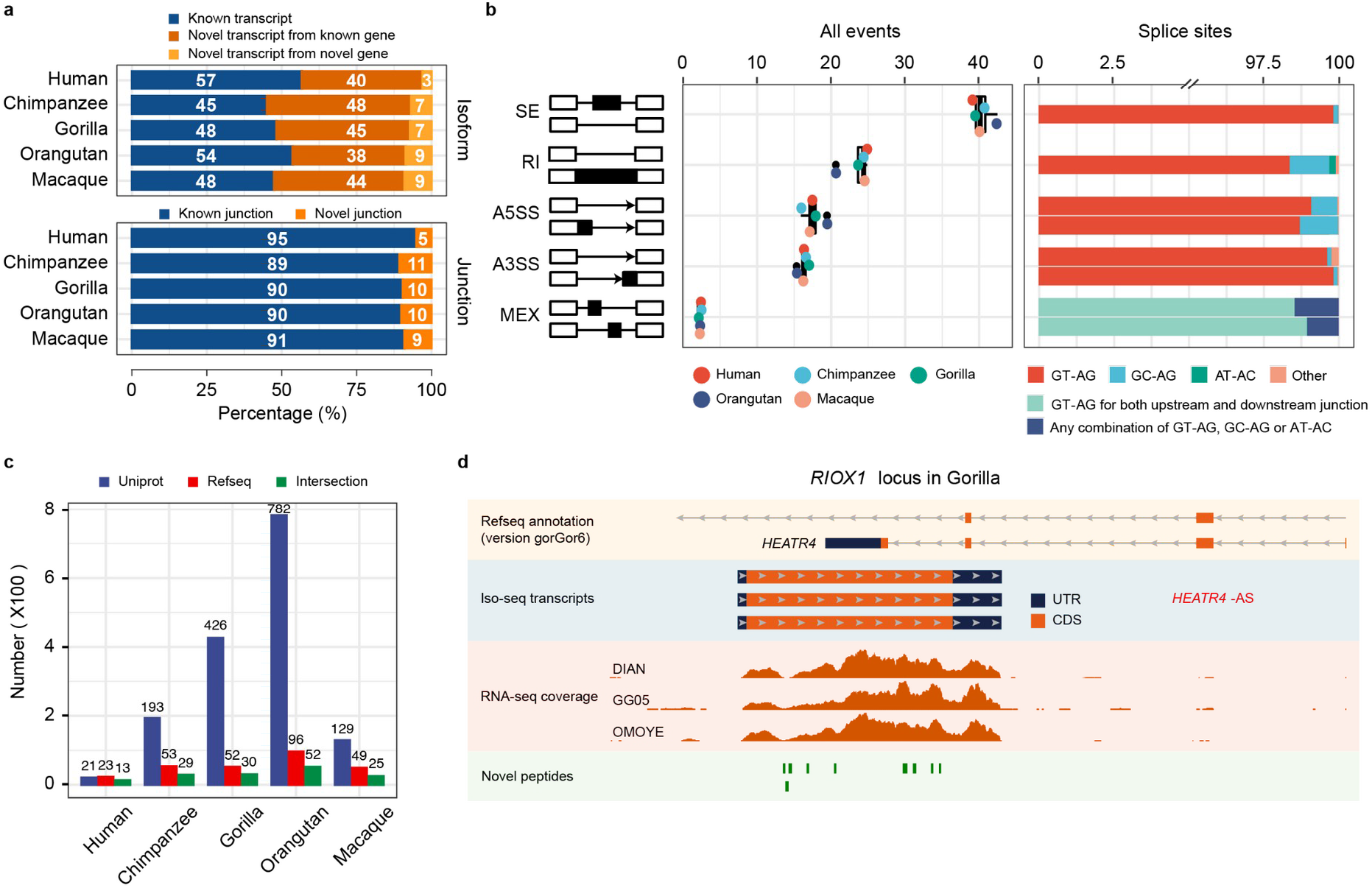
Isoform sequencing and mass spectrometry proteomics largely improve reference annotations. **a)** Percentage of known *versus* novel transcripts (top) and junctions (bottom) captured by Iso-seq (Ensembl V91). **b)** Percentage of alternative splicing (AS) events detected by Iso-seq (left) and percentage of splice site combinations associated with each AS class (right). SE, skipping exons; RI, retained introns; A5SS, alternative 5’ splice sites; A3SS, alternative 3’ splice sites; MEX, mutually exclusive exons. **c)** Number of annotated genes with novel detected peptides in each species for UniProt reference proteomes, NCBI RefSeq CDS annotations, or both. UniProt annotations are based on hg38, panTro5, gorGor4, ponAbe2, and rheMac8. NCBI RefSeq CDS annotations are based on hg38, panTro6, gorGor6, ponAbe3, and rheMac10. Peptides mapping to multiple isoforms are included. **d)** Example of a novel gene (*RIOX1 locus* shown in gorGor6 assembly) supported by Iso-seq, RNA-seq, and mass spectrometry in gorilla samples.

### Mass spectrometry detects missing proteins in NHP reference proteomes

Since most Iso-seq transcripts are predicted to be protein-coding, we performed high-throughput mass spectrometry experiments to search for evidence of peptide translation. To do so, we predicted the ORF of all Iso-seq transcripts in all species and built a comprehensive protein database that constitutes our search space for peptides across human and NHP LCLs (Methods). Within this search space, we detect a total of 49,428 distinct tryptic peptides (FDR<5% and Mascot Ion Score>20, excluding contaminants), from which over 24,900 are confidently detected, on average, in each species (reporter ion intensity signal (abundance) per sample > 50 in any sample of a given species, and species-wise median ratio of sample abundance to pool abundance > 0.6, see Supplementary Table 4, Supplementary Data 2 and Methods). These detected peptides are more frequently annotated in reference proteomes (>99%) than all theoretically predicted peptides from Iso-seq transcripts (~87%).

A number of Iso-seq derived peptides detected in each species are novel in the corresponding UniProt reference proteomes (Methods), including 22 for human (hg38), 320 for chimpanzee (panTro5), 882 for gorilla (gorGor4), 1,590 for orangutan (ponAbe2), and 207 for rhesus macaque (rheMac8) (see Fig. 2c for the number of annotated genes with novel detected peptides). To explore these results in the context of the most recent primates assemblies, we then searched the detected peptides in the new CDS annotations (NCBI RefSeq CDS), which provide not only curated proteins (NP) but also *in silico* predictions (XP). On average, nearly 78% of our detected tryptic peptides in NHP are only present in predicted entries (XP) from NHP RefSeq proteomes, highlighting the value of our multi-species mass spectrometry experiments in primates to confirm such protein predictions for the first time. When considering both curated and predicted proteins, we detected 25 novel peptides in hg38, 98 in panTro6, 94 in gorGor6, 225 in ponAbe3, and 75 in rheMac10 RefSeq CDS sets corresponding to novel and annotated genes (Fig. 2c and Supplementary Data 3). These numbers clearly illustrate the distinct levels of incompleteness in primate reference proteomes, reflecting the differences in the previous efforts made by the community to complete the gene repertoire and study the proteome of each species.

These improvements in proteome annotations provide a vantage point to study the expression of proteins in primates. Unannotated genes with multiple supportive peptides were also detected in NHP (Supplementary Data 3), as for *RIOX1* in gorilla samples (Fig. 2d), which is classified as a pseudogene in Ensembl (gorGor4) and absent in RefSeq (gorGor6) despite its importance in cancer research ^24^. In other cases, we expand the exon and CDS annotations, as in the gene *SON* in orangutan (ponAbe3), a key splicing factor in cell-cycle progression involved in the response to hepatitis B virus ^25–28^ (Supplementary Fig. 4). Our results demonstrate that integrating cross-species Iso-seq with mass spectrometry significantly improves proteomic annotations even in the most recent NHP assemblies.

### Emergence of species-specific genomic regions under active transcription

To study the recent transcriptome evolution in primates, we first focused on the genomic presence/absence of the expressed transcripts in the different species. To do so, we inspected orthologous transcripts produced by one-to-one orthologous genes in the five genomes that are expressed in at least one species (Supplementary Fig. 5, Methods). A set of 49,797 transcript models from 7,858 genes was projected to the five primates highlighting the high genomic conservation of the transcript structures (Supplementary Data 4 and 5). In addition to this high conservation, we found 61 species-specific exons without an orthologous counterpart in the other primates (Supplementary Data 6, Methods). A significant fraction of them (72%) have not been annotated before, which is possibly due to their lower expression levels (Supplementary Fig. 6).

These species-specific exons are frequently located in repetitive regions (79% of them overlap repeat elements), particularly SINEs (mainly Alu elements) and LINEs (Supplementary Fig. 7), which are associated with exonization events ^29^. We then wondered if these exonizations significantly alter the CDS of the encoded protein. We found that the incorporation of these exons in the transcript usually preserves the combination of protein domains (96.7% of all exonizations), encoding protein variants with likely conserved functions. In other cases, the new exon is included in untranslated regions (UTR). For instance, in our LCLs, the exonization of three repeat elements in human *MRNIP* results in a longer 3'UTR with predicted small interfering RNA (siRNA) binding sites (Supplementary Fig. 8), the Alu exonization in gorilla *NPLOC*4 lengthens the encoded protein by 38 amino acids, and the exonization of an Alu element in *ABCD4* generates a new open reading frame in orangutan, preserving all *ABCD4* protein domains.

### Isoform innovations revealed by single-molecule sequencing

The high conservation of the gene structures led us to analyze the evolution of transcript expression in primates. We focused on the set of orthologous transcript models in the five genomes, and quantified transcript expression levels using high-depth RNA-seq data from 15 LCLs (3 LCLs per species) to obtain accurate expression estimates (Methods, Supplementary Data 7). Consistent with widespread transcriptome conservation, a high number of transcripts are expressed in all five species (N=18,325 transcripts), while 55% of all projected transcripts are only expressed in some species. Thus, to further understand the patterns of transcript expression gains and losses in the primate lineage, we reconstructed the evolutionary dynamics under Wagner parsimony ^30^ and uncovered thousands of transcripts whose expression has been gained or lost across primates. We reasoned that at least part of the transcripts not detected in all species can result from inter-individual transcript expression variability. To account for this variability, we established stringent criteria to keep the innovations that are consistent across all biological replicates (Fig. 3a and 3b, Supplementary Data 8, Methods). In fact, these criteria increase the percentage of transcript gains and losses that are coherent with the phylogeny (*i.e.*, being gained or lost just once in the phylogeny) in comparison to the total set of transcripts (Supplementary Fig. 9).

**Figure 3.**
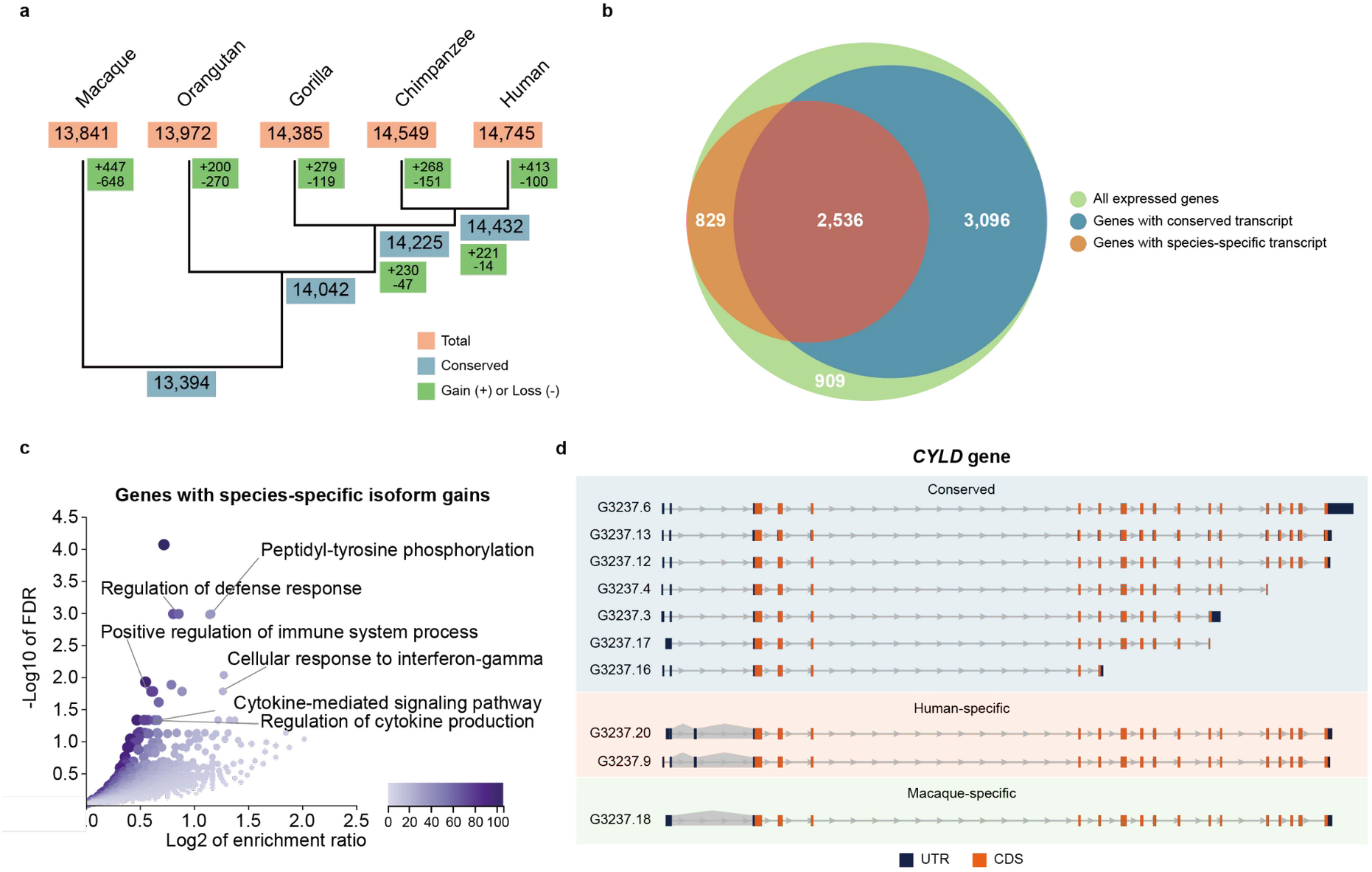
Transcript expression gains and losses in the primate lineage. **a)** Dynamics of transcript expression gains and losses in 5 primate species (inferred by Wagner parsimony). The phylogenetic tree is built based on the known phylogeny. Transcript expression gains and losses in macaque *versus* great apes are shown in macaque’s branch, although the direction (gain or loss in macaque or great apes) cannot be determined. **b)** Overlap of genes expressing highly conserved (blue) and species-specific transcripts (orange). **c)** Functional enrichment for genes displaying species-specific isoform innovations (N=868 genes). Over-representation analysis (ORA) was conducted using the Gene Ontology Biological Process database (affinity propagation clustering). False discovery rates (FDR) were adjusted using the Benjamini-Hochberg method ^53^. Color legend indicates the number of overlapping genes between the gene set under evaluation and the pathway gene set. **d)** Example of a gene, *CYLD*, expressing transcripts in all 5 species (Conserved, blue shadow), exclusive expression in human (Human-specific, orange shadow), and exclusive expression in rhesus macaque (Macaque-specific, green shadow). Grey shadows indicate species-specific splice junctions. Untranslated regions (UTR) and coding sequences (CDS) are colored in blue and orange, respectively.

We defined 1,215 high-confidence species-specific transcripts (only expressed in a single species). The vast majority of them (94%) arise from species-specific splice junctions, with exon skipping being the main mechanism involved in these innovations (Supplementary Fig. 10). Furthermore, we confirmed that although gene expression upregulation might result in species-specific transcript detection, our detections of species-specific gains are not driven by increased gene expression levels in the corresponding species (90.21% of genes expressing species-specific transcripts are not significantly upregulated in the corresponding species, see Methods). These species-specific transcripts are enriched in annotation novelties (OR=3.86, Fisher’s exact test P < 2.2e-16), demonstrating the suitability of our strategy to uncover primate isoform innovations otherwise missed by reference transcriptomes.

Importantly, the genes displaying species-specific innovations show an over-representation in innate immune system functions (Fig. 3c, Supplementary Table 5, Methods), and are more often involved in immune system-related diseases (*e.g., OAS2*, *OAS3*, *RNASEL*, *IRF7* and *JAK1;* Fisher’s exact test P=0.016, OR=1.50). Given the involvement of these genes in the innate immune response, we then asked if they were enriched in any functional subgroup according to a curated classification of innate immunity genes ^31^. Notably, immune sensors (Fisher’s exact test P=0.019, OR=1.96), effectors (P=0.007, OR=2.5), and secondary receptors (P=0.027, OR=3.31) are more likely to display species-specific isoform innovations in primates. Furthermore, genes expressing species-specific transcripts tend to be highly expressed in the spleen, which actively participates in the immune response (GTEx expression data across all human tissues, Hypergeometric test −log10 corrected P=3.14, FC=1.95) (Methods). We also found that these genes expressing species-specific isoform innovations are enriched in genes involved in cell proliferation, consistent with an enrichment in LCL-specific functions (Fisher’s exact test P=0.002, OR=1.61). In contrast, we observed that this set of genes is depleted in housekeeping genes (Fisher’s exact test P=5.46e-06, OR=0.64), showing that the essential role of housekeeping genes is also reflected in the conservation of their global splicing structures.

An illustrative example of transcriptomic innovation is *CYLD*, a tumor suppressor deubiquitinase involved in NF-κB activation ^32,33^, whose splicing patterns impact the regulation of immune cells activation ^34,35^. In LCLs, we observe species-specific expression in some *CYLD* isoforms (Fig. 3d). In this case, the inclusion of a non-coding exon generates two human-specific transcripts, while an alternative acceptor site produces a macaque-specific transcript. Another example is *IFI44L* (interferon-induced protein 44-like), which is involved in the immune response to infection and autoimmune disorders ^36–46^ and whose isoform expression has also been found to be clinically relevant ^47,48^. *IFI44L* expresses three human-specific transcripts encoding a different ORF than the isoforms expressed in all species (Supplementary Fig. 11). Our results indicate that immune system and cell proliferation-related genes are preferentially targeted by recent transcriptomic innovations within the 7,858 orthologous genes expressed in LCLs.

### The impact of *cis*-regulatory genetic changes in isoform innovation

The efficient inclusion of exons into mature transcripts is determined mainly by splicing signals located in the exon boundaries. Thus, interspecies genetic changes affecting the orthologous splice sites might contribute to events of transcript diversification, including the emergence of species-specific isoforms. In the context of our transcriptomes, we examined the sequence conservation of orthologous splice sites (terminal intron dinucleotides). Overall, we observed widespread sequence conservation of splice sites across primates, reflecting strong selective constraints in these regions (Supplementary Fig. 12). To explore high-impact interspecies changes in splice sites, we then classified all orthologous junctions into canonical and non-canonical, associated with high and low splicing efficiencies, respectively. After filtering out lowly expressed isoforms and genes (Methods), we observed 82,527 splice junctions present in our transcriptomes, from which 1,318 show a change in canonicity in any species. This scenario affects 16% of all evaluated genes (N=1,008 genes), which are enriched in T (FDR=0.019) and B-cell activation (FDR=0.019) as well as JAK/STAT signaling (FDR=0.022) pathways (Supplementary Table 6, Methods), showing that *cis*-regulatory changes in splice sites are primarily associated with genes involved in immune system functions.

We then investigated the contribution of these genetic changes to the expression of species-specific isoforms. We observed that 69% of the high-confidence isoform innovations (833 out of 1,215 isoforms) harbor conserved canonicity of their splice sites in all species. Remarkably, the expression of 184 transcripts is driven by genetic differences producing canonical splice sites only in the species in which they are expressed. The enrichment in immune system functions holds for species-specific transcripts regardless of if they occur with highly conserved orthologous splice sites or not, showing the global relevance of isoform innovation in the evolution of the immune system (Supplementary Fig. 13).

An example of interspecies genetic differences in the splice sites is *RRP15*, a gene involved in cell proliferation and apoptosis ^49^. In *RRP15*, a high-frequency variant only present in modern humans ^50^ disrupts an acceptor site of a coding exon (from AG to TG), and a downstream splice site is used instead. This mutation results in a human-specific exon shortening representing a deletion of three amino acids (disordered residues involved in protein binding), whereas the reading frame is preserved compared to other primates ^51,52^ (Supplementary Fig. 14). In other scenarios, the genetic change in splice sites promotes the transcription of a new exon, as we observe in gorilla LCLs for *NAP1L4* gene (Supplementary Fig. 15).

### Evolutionary dynamics of isoform usage patterns in the primate lineage

After studying the evolution of switching on/off isoform transcription, we aimed to investigate the evolution of isoform transcription levels. To disentangle isoform-specific transcription regulation from changes in total gene expression, we assessed the relative transcript usage across the five species. To do so, we computed sample-wise fractions of isoform usage (IU) in our quantified orthologous transcripts (Methods). Principal component analysis and hierarchical clustering indicate that IU values cluster by species and that differences across species increase with evolutionary distance, reflecting the known phylogeny (Fig. 4a, Methods). As expected, IU values are higher in annotated transcripts than in novel isoforms (Welch Two Sample t-test P < 2.2e-16), indicating that isoforms displaying important contributions to the expression of their genes are better represented in reference annotations.

**Figure 4.**
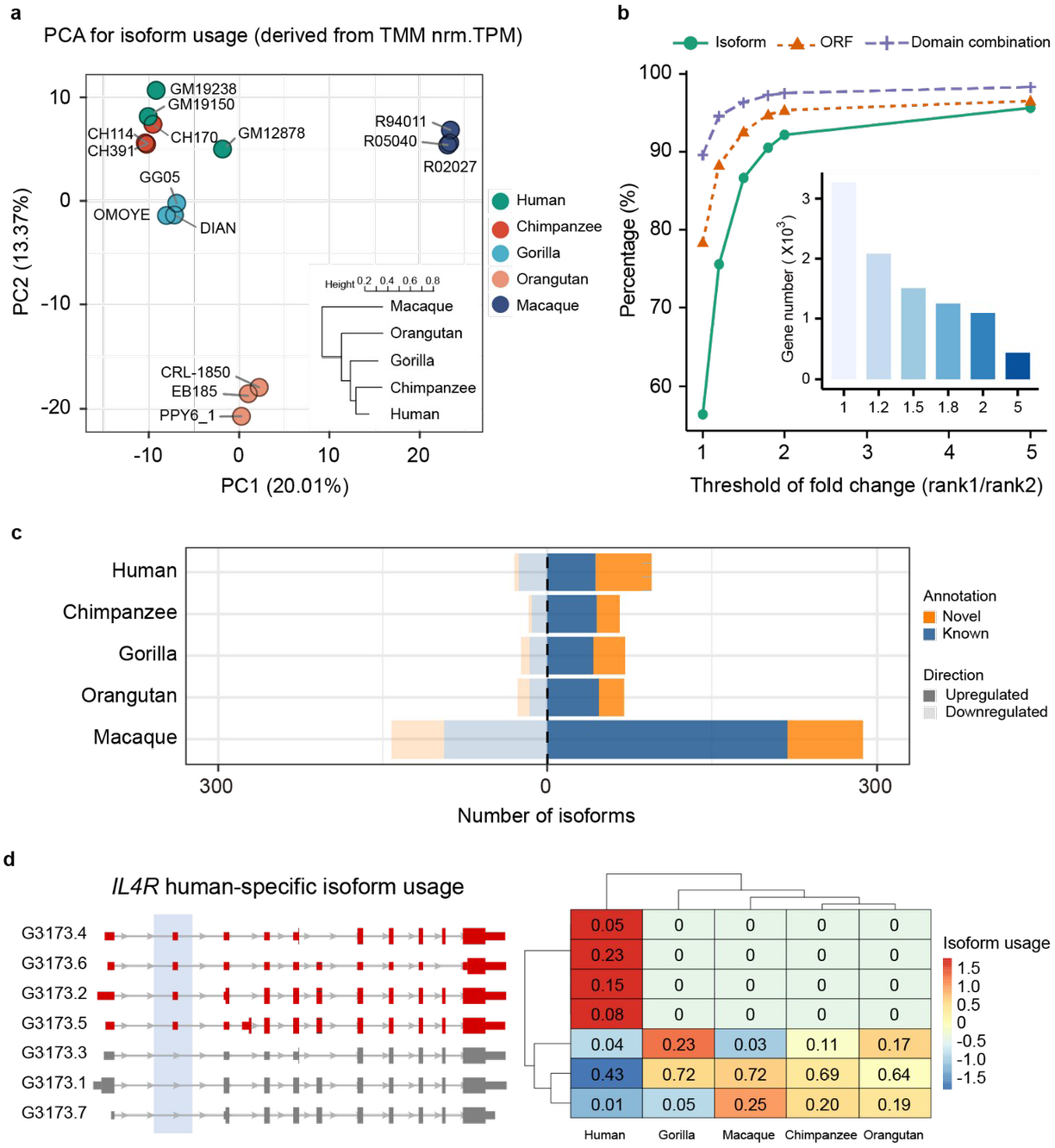
Isoform usage dynamics across primates. **a)** Principal component analysis (PCA) for isoform fraction values (IU) of 43,484 transcripts from one-to-one orthologous genes. A hierarchical clustering for the euclidean distances of IU Spearman correlations across samples is also shown. IU values were calculated from Transcripts Per Million (TPM) after batch effect correction (comBat) and TMM (Trimmed Mean of M-values) normalization (Methods). **b)** Percentage of rank 1 isoform (green), ORF (orange), and protein domain combination (purple) shared by all primates by different thresholds of expression fold change between rank 1 and rank 2 isoforms. The number of genes considered for each threshold is indicated in the inner bar plot. **c)** Number of isoforms showing species-specific usage changes. Upregulated (dark shadow) and downregulated (light shadow) isoforms are classified into novel (orange) and known (blue) isoforms based on the reference transcriptome in the corresponding species (NHP novel assemblies). **d)** *IL4R* human-specific isoform usage. Human-specific upregulated isoforms are colored in red, and non-differentially used isoforms in grey (left). Isoform usage (IU) values across species are shown in the heatmap (right), where colors represent row-scaled IU values (legend).

To investigate the usage conservation of transcripts with a major contribution to the total gene abundances, we then focused on the isoforms with the highest IU across primates (rank 1 and rank 2 transcripts) after filtering out genes with very similar transcripts (internal exon extensions leading to quantification ambiguities, see Methods). Since different isoforms can produce identical proteins, we also evaluated the conservation of the ORF and of the combination of protein domains encoded by the dominant transcripts (rank 1) in different species. Notably, all species share the same dominant transcript for 55.44% of the genes, a trend that is more evident as bigger fold changes of expression between rank 1 and rank 2 transcripts are considered (Fig. 4b), suggesting that changes in the dominant transcript are less common. This conservation is higher for the protein-coding features, with 78.33% and 90.01% of genes sharing identical ORF or combinations of protein domains in their dominant transcripts in all species, respectively, consistent with changes in the dominant transcript being almost always functionally conservative. Our results highlight the greater constraints associated with protein structure and function conservation, whereas distinct preferences in isoform UTR or even subtle ORF changes are more frequent in the dominant transcripts across the primate phylogeny.

We investigated other changes of relative transcript usage across primates. To do so, we conducted pairwise comparatives to find interspecies differential isoform usage (DIU, Methods) after excluding lowly expressed genes and isoforms ^54^ as well as genes expressing highly similar transcripts (variability in internal exon boundaries) (Methods). In a scenario of broadly conserved isoform usage across species, we detected over 800 transcripts showing species-specific usage changes (Supplementary Data 9), a significant fraction of which are missed by reference annotations (Fig. 4c). LCL-specific genes display species-specific DIU changes more frequently (Fisher’s exact test P=0.015, OR=1.69), although we did not observe any pathway enrichment for the genes displaying species-specific DIU. An example of human-specific differential isoform usage is the interleukin-4 receptor (*IL4R*), a gene under positive selection in primates ^55^, which plays a major role in the immune response ^56^. We detected four human-upregulated *IL4R* transcripts, two of them encoding shorter *IL4R* proteins (Fig. 4d).

### Local differential splicing dynamics in transcribed regions

While full-length isoform expression levels provide an accurate view of major usage changes in clearly different transcripts, other local splicing changes are better detected by total exon expression levels that accumulate the signal of all transcripts including the exon. We assessed the differential exon usage (DEU) across the primate phylogeny to obtain a comprehensive view of isoform transcription evolution (Methods). We first leveraged our projected Iso-seq models to define 201,191 orthologous exonic parts (*i.e.,* transcribed regions) from 7,766 one-to-one orthologous genes. Then, RNA-seq read counts in these exonic parts were obtained across all samples to detect interspecies differences in relative exon usage (Methods). We observed nearly 4,000 exonic parts displaying species-specific local usage changes (Supplementary Data 10), with skipping exons (SE) being a prevalent AS event among them (see Fig. 5a for a description of features in SE showing species-specific usage changes). As for the above reported exons that are only present in the genome of a single species, we observed significant enrichment of Alu elements in human-specific upregulated exonic parts compared to downregulated or regions of conserved usage (Supplementary Fig. 16). This suggests that Alu insertions contribute to increasing orthologous exon inclusion frequencies, especially for the case of SE. This is consistent with previous reports on the Alu-mediated modulation of exon usage rates ^57^.

**Figure 5.**
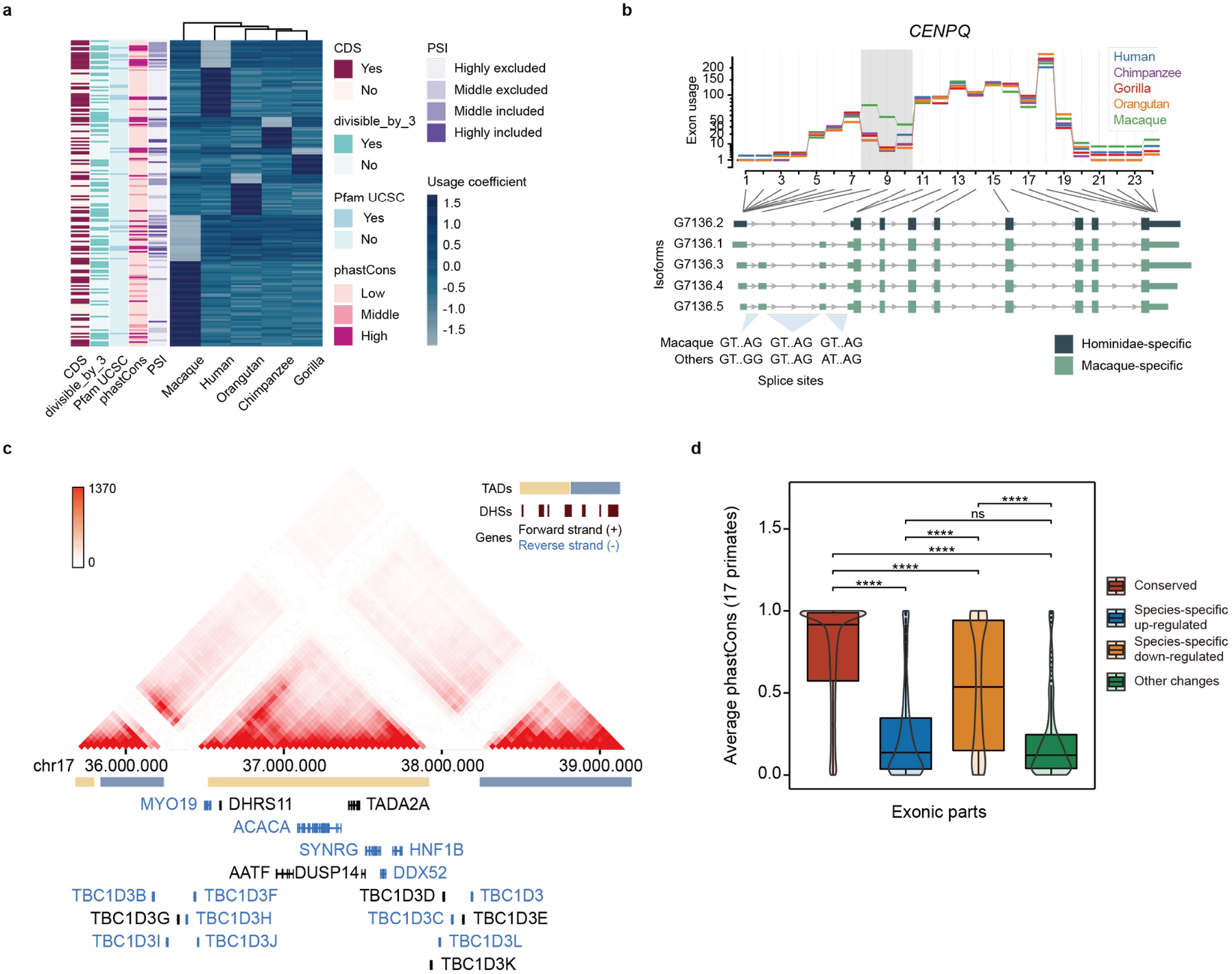
Exon usage dynamics across primates. **a)** Usage coefficients for skipping exons displaying species-specific changes (row-wise scaled). Overlap with coding sequence (CDS) and Pfam protein domains (UCSC), divisibility by 3, average percent-splice-in (PSI) and average phastCons scores are shown for each exon (see Methods). **b)** Example of gene (*CENPQ*) with different splice sites in macaque and great apes. Exon usage coefficients are illustrated at the top. Hominidae+ and macaque-specific Iso-seq transcripts are color-coded (bottom). **c)** chr17q12 region with high density of human-specific DEU. Hi-C interaction intensity is indicated in the color legend. Gene orientation, DNase hypersensitive sites (DHS) and topologically associating domains (TADs) are also displayed. Average conservation score in primates for exonic parts with conserved usage (N=95,453), species-specific upregulation (N=941) or downregulation (N=1,340), and exonic parts showing other usage changes (N=277) (Methods). Exonic parts longer than 5 bases (all included as passed in 1000 Genomes hg38 strict mask) are shown. Statistical significance of the difference across groups was assessed by Wilcoxon test and adjusted by Holm method (**** = P ≤ 0.0001, ns = P > 0.05).

Notably, genes with species-specific DEU are enriched in functions related to the regulation of immunity, such as interferon signaling and innate immune response (*IRF2*, *IRF3*, *IRF5*, *IRF7*, *IRF9*, *IFI35*, *IFIT1*, *ISG20, IFNGR2, HLA-DQB1, CIITA,* etc.), as well as proliferative (DNA repair, centrosome localization, etc.), housekeeping (ncRNA processing, mitochondrial gene expression, etc.), and viral life-cycle control functions (FDR ≤ 0.05, Supplementary Table 7 and Methods). An example of this is *CENPQ*, a centromere protein involved in autoimmune disease and mitosis progression ^58,59^ whose species-specific DEU is driven by a genetic change in the primate splice sites. In macaque, two junctions harbor GT-AG splice sites, leading to the recognition of two 5’ UTR exons and the preferred usage of the isoforms that include them. On the contrary, great apes -where these splice sites are not canonical- only express the transcript that skips these exons (Fig. 5b). We also noticed that 13.89% of genes displaying species-specific DEU co-occur with gene expression up or downregulation in the same species. Six genes downregulated in human LCLs (*SYNRG*, *DUSP14*, *TADA2A*, *ACACA*, *MYO19,* and *DDX52*) are comprised in the same topologically associating domain in chr17q12 ^60^, and display human-specific exon usage changes (Fig. 5c). Genomic rearrangements in this locus have been extensively studied given their evolutionary relevance and implications in many diseases, including neurodevelopmental disorders ^61–64^. To date, this is the first report of human-specific splicing changes occurring in multiple genes encoded in this region.

We found that exonic parts experiencing species-specific usage upregulation display lower conservation in primates than downregulated exonic parts in a given species (Fig. 5d). This higher variation in species-specific upregulated regions is also reflected in the nucleotide diversity within human populations (see Methods, Supplementary Fig. 17). When focusing on transcribed regions with human-specific usage changes, we noticed upregulated exonic parts are depleted in CDS (Supplementary Fig. 18) and are less present in principal isoforms (defined by APPRIS ^65^) compared to human-specific downregulated or regions of conserved usage (Supplementary Fig. 19). Our findings suggest that species-specific gains in exon usage are associated with fast-evolving regions whose changes in usage lead to subtle or regulatory changes in the main function of these genes.

We reasoned that genes with splicing changes detected across primates might show characteristic selective pressures in their coding sequences. To investigate this possibility, we assessed the ratios of nonsynonymous to synonymous substitutions (*d*N/*d*S) in human populations for the genes under evaluation ^55^. To do so, we combined our results from isoform gains and losses, DEU and DIU analyses to define gene classes according to their splicing patterns: those showing any species-specific upregulation signal in human (N=278 genes) or NHP (N=1,239 genes), those displaying multiple events of species-specific upregulation in several species (N=287 genes), those experiencing other splicing changes consistent with differences between groups of species (N=1,174 genes), and genes with fully conserved splicing patterns (N=4,532 genes) (Methods). Remarkably, we observed significantly higher *d*N/*d*S in human populations for the gene sets showing splicing changes involving NHP than for genes with fully conserved splicing, while the genes with only human-specific upregulation show intermediate *d*N/*d*S (Supplementary Fig. 20). These results suggest a link between dynamic splicing evolution in primates and accelerated or less constrained coding changes in human populations, with intermediate selection constraints in genes with only human-specific upregulation.

To further investigate the occurrence of interspecies splicing changes in genes under adaptive selection, we then investigated if the above-defined gene classes showing splicing changes were enriched in signals of positive selection in primates ^55^ in comparison to genes with conserved splicing patterns. We found that only genes showing independent events of species-specific upregulation in multiple primates are significantly enriched in signals of positive selection (Fisher’s exact test, Bonferroni adjusted P=0.014, OR=2.90), indicating that they might be important targets of adaptive evolution. This set of genes accumulating recurrent species-specific changes in the splicing programs of different primates is enriched in genes associated with the regulation of the immune response (Gene Ontology Biological Process ORA, FDR=0.03, enrichment ratio=2.17), reinforcing the link between evolutionary changes in splicing and the immune function.

## Discussion

To date, our knowledge of the evolution of primate transcriptomes is very limited. Here, we have produced the most extensive catalog of full-length isoforms across the primate lineage complemented with matching Illumina RNA-seq data. The combination of both allowed us to create highly curated transcriptomes consisting of a high proportion of novel transcripts from annotated genes as well as a number of antisense and intergenic transcripts. Furthermore, new AS events have been identified, and we have observed a conserved composition of AS classes in great apes and rhesus macaque. This resource greatly expands the available gene annotations for model and non-model primates by providing a more accurate and complete view of genes and isoforms, which are crucial for further genomic and functional studies.

We have taken advantage of our curated isoform catalog to predict the corresponding ORF for coding transcripts, representing the majority of the sequenced transcriptomes. This ORF collection has been fundamental to build proteomic databases assisting coupled mass spectrometry analysis. In this way, we detected evidence of translation for Iso-seq transcripts identifying peptides that are not present in reference proteomes. We detected fewer novelties at the proteome than at the transcriptome level, reflecting a combination of the differences in experimental sensitivities, in the completeness of proteome and transcriptome annotations, and in the proteome and transcriptome complexities ^66^. All considered, our analyses show that there is still room for improvement of proteome annotations by integrating isoform resolution sequencing data.

Our results show a significant number of transcript expression gains and losses in a scenario of widespread conservation of transcript expression in the primate lineage. We found that genes involved in the regulation of innate immunity are enriched in isoform innovations even when controlling for the composition of LCLs transcriptomes. From an evolutionary point of view, many immunity-related genes have undergone strong selective pressures in response to constantly evolving host-pathogen interactions, leading to the accelerated evolution of these genes ^67,68^. However, very few studies have addressed how these rapid evolutionary rates are reflected in the immune system-related splicing programs in the context of recent human evolution ^69,70^. On the other hand, exon skipping events are enriched in species-specific transcript gains, posing this type of alternative splicing as a prevalent mechanism in transcriptomic innovation. We detected a number of high-impact genetic changes located in orthologous splice sites, which are clearly involved in shaping the splicing preferences in different primate species. We have also characterized the emergence of new exons, many of them absent in reference annotations, and confirmed the association between the exonization process and the insertion of transposable elements such as SINEs and LINEs.

The patterns of isoform usage recapitulate the phylogenetic structure in primates, with the most abundant transcripts of each gene frequently encoding the same ORF or showing conservative changes across species. This also highlights that the evolution of recent isoform transcriptional changes usually preserves the functional integrity of the protein encoded by the gene, pointing to these changes as a source of subtle regulatory fine-tuning for important genes. Here, we report a number of interesting species-specific changes in isoform preferences, involving a significant fraction of isoforms missed by reference transcriptomes. Further analyses will be required to understand the functional implications of the splicing changes affecting coding and non-coding regions. In this direction, it is remarkable that our set of differentially used exons is enriched in untranslated regions, which harbor binding motifs for RNA-binding proteins and small RNA molecules (*e.g.*, miRNAs), playing fundamental roles in the regulation of isoform stability and localization ^71^. Furthermore, the enrichment in signals of positive selection in the coding sequences of genes accumulating independent species-specific gains of isoform expression, isoform usage, or exon usage in multiple primates suggests a link between recurrent splicing changes and adaptive evolution currently occurring in extant primates.

In summary, this study expands our current knowledge on transcript and protein diversity in human and non-human primates providing a comprehensive evaluation of the dynamics of transcriptome evolution in the primate lineage as reflected in B-cells-derived LCLs. The combination of high-depth long and short read transcriptomics and high-throughput mass spectrometry provides an excellent framework to characterize many evolutionary novelties missed by available transcriptome and proteome annotations. The discovery of these recent innovations highlights the prevalence of splicing changes during primate evolution. Our results show that genes involved in the immune response, in cell proliferation and with cell-type specific expression profiles have been the target of numerous transcriptional innovations. The strong adaptive demands of the immune response and cell proliferation at the transcriptional level are satisfied mostly by incorporating fast regulatory and subtle changes without drastically impacting the viability of the response. Moreover, this set of evolutionary changes constitutes the basis for future functional assays for studying their role in interspecies phenotypic divergences in the immune response and cell proliferation, areas of major interest in evolutionary and biomedical research.

## Methods

### Lymphoblastoid cell lines origin and culture

Lymphoblastoid cell lines were kindly provided by Dr. Antoine Blancher (chimpanzee LCLs CH507, CH170 and CH322), Dr. Aurora Ruiz (gorilla LCL GG05), Dr. Chris Tyler-Smith (orangutan LCL CRL-1850) and Dr. Gaby Dioxiadis (macaque LCL R94011). GM19238 (human) and EB185(JC) (orangutan) were purchased from Coriell Institute and Sigma, respectively. The remaining cell lines used in this article are described in ^72^.

Cells were cultured in RMPI-1640 with L-Glu (Gibco) supplemented with 15% FBS (Gibco) and 1% P/ S (10000 U/mL, Gibco). Multiple cell pellets totaling 12×10^6^ cells per cell line were stored at −80°C until further extraction after 2 washes with DPBS.

### DNA and RNA extraction

A pellet consisting of 2×10^6^ cells was washed once and resuspended in DPBS to use as starting material to extract DNA using the DNeasy Blood and Tissue Kit (Qiagen) following the manufacturer’s instructions. RNA was extracted from a 3×10^6^ cell pellet, previously stored in QIAzol Lysis Reagent, using the miRNeasy mini kit (Qiagen) and following the manufacturer’s instructions.

### RNA-seq data production and processing

Illumina TruSeq Stranded RNA-seq experiments were conducted at Macrogen for the following LCLs: GG05, CRL-1850, GM19238, CH507, CH170, CH322, EB185 and R94011, and complemented with the RNA-seq data described in ^72^. RNA-seq data from GM12878 was retrieved from NCBI (accession SRR998197 and SRR998198). RNA-seq reads were filtered with SOAPnuke ^73^ after adapter removal. Filtered reads were first aligned to each primate genome with STAR ^74^ to scan for supported splice junctions and quantify gene, transcript and exon expression levels.

### Isoform sequencing data production

Two lymphoblastoid cell lines (LCLs) per species were used for isoform sequencing experiments (PacBio Iso-seq) (human: GM12878, GM19150; chimpanzee: CH114, CH391; gorilla: DIAN, OMOYE; orangutan: two isogenic cultures of PPY6; macaque: R05040, R02027). In the first run, three libraries of 1-2 kb, 2-3 kb, and >3 kb were generated using SMRTbell Template Prep Kits and sequenced on a PacBio Sequel system at Novogene Co. Ltd. A total of 26 Gb (subread data) was generated per species, with an average length and N50 of 1,980 bp and 2,399 bp, respectively. In a second run, libraries were produced without size selection using SMRTbell Template Prep Kits and sequenced on a PacBio Sequel system. A total of 56 Gb (subread data) was generated per species, with an average length and N50 of 2,162 bp and 2,647 bp, respectively.

### Iso-seq data processing

Size-selected PacBio Sequel subreads were collapsed into circular consensus sequences (CCS), and only sequences with 5’ and 3’ primers and poly-A tail were selected for further analyses (full-length non-chimeric CCS; FLNC) following the IsoSeq3 workflow (PacBio SMRTAnalysis software). Then, FLNC sequences were clustered by Iterative Clustering and Error Correction (ICE) and corrected by Arrow algorithm, generating a set of full-length high-quality polished isoforms per species.

Hybrid correction of full-length high-quality polished isoforms was carried out by LoRDEC ^75^ using sample-specific paired-end Illumina RNA-seq data. Three matched RNA-seq datasets for each LCL were used except for GM12878, for which two publicly available datasets were used (NCBI accession SRR998197 and SRR998198).

After poly-A trimming (trim_isoseq_polyA script from official PacBio Github repository), Iso-seq isoforms were mapped to the reference genomes hg38 (human), panTro5 (chimpanzee), gorGor4 (gorilla), ponAbe2 (orangutan), and rheMac8 (rhesus macaque) using GMAP ^76^. Illumina RNA-seq data was used to further attenuate the errors derived from Iso-seq and refine the transcript splice junctions ^77^.

Uniquely mapped isoforms were used as input for Cupcake ToFU scripts (https://github.com/Magdoll/cDNA_Cupcake). At this moment, low-quality alignments were discarded based on identity (95%) and coverage (99%), and isoforms were collapsed into unique transcripts. This step removes the remaining redundancy accounting for 5’ end variability since ICE performs a very conservative clustering.

### Artifact filtering from Iso-seq data

Transcript isoform FASTA files were provided to SQANTI ^23^ for quality control and artifact filtering. Splice junction coverage by RNA-seq and isoform abundances (full-length reads and TPM) were used to train the SQANTI classifier. Splice junction coverage in Iso-seq isoforms was obtained by mapping sample-specific RNA-seq to the corresponding reference genome with STAR ^74^. Normalized quantification estimates for Iso-seq isoforms (TPM) were computed using Kallisto-quant ^78^ with --rf-stranded argument. Finally, the SQANTI machine learning method (sqanti_filter.py) was applied to the collapsed isoforms in each species to keep a set of highly confident isoforms in each species. To be considered novel, an isoform must have a splice junction structure that is not annotated in the corresponding reference transcriptome. Data-manipulation was performed using SAMtools ^79^, BEDtools ^80^, and in-house scripts.

### Classification of alternative splicing patterns

A classification of alternative splicing (AS) events was generated by SUPPA ^81^ from the GTF files resulting from SQANTI. Five types of AS were identified: skipping exon (SE), alternative 5’ or 3’ splice sites (A5SS/A3SS), retained introns (RI), and mutually exclusive exons (MX). AS genes were defined as genes with any associated AS event. Splicing events were considered as known if both splice events (inclusion and exclusion forms of the AS event) are found in Ensembl V91 transcriptomic annotations.

### Protein extraction and precipitation

Two million cell pellets were thawed on ice, washed within 1ml of cold DPBS, and further centrifuged at 2500 × *g* 5 minutes at 4°C. After removing the supernatants, 450 μl of cold RIPA buffer (Thermo Scientific) supplemented with 10μl/ml Halt™ Protease and Phosphatase Inhibitor Cocktail (100X) (Thermo Scientific) was added to disrupt cell pellets. After incubating 5 minutes the tubes swirling on ice, the suspension was sonicated for 30 seconds with 50% pulse a total of 3 times with 10 seconds of rest using the Branson 250 sonicator. Later, tubes were further incubated for 30 minutes on ice swirling. Finally, tubes were centrifuged at 14000 × *g* for 15 minutes at 4°C to pellet the cell debris, and supernatants were transferred into new protein LoBind tubes (Eppendorf). Extracted proteins were quantified using Pierce BCA Protein Assay Kit (Thermo Fisher Scientific) and stored at −80°C until further analysis.

To precipitate the proteins, 6 volumes of overnight chilled acetone (−20°C) were added, then tubes were inverted 3 times and incubated overnight at −20°C. The next day tubes were centrifuged at 16000 × g for 10 minutes at 4°C, and acetone was removed carefully without disturbing the protein pellet.

### Mass spectrometry sample preparation

Three biological replicates for each species were processed. Samples (12-120 μg) were solubilized in 100mM triethylammonium bicarbonate (TEAB) at a concentration of 1 μg/μl, reduced with tris(2-carboxyethyl)phosphine (0.12-1.20 μmol, 37°C, 60 min) and alkylated in the dark with iodoacetamide (0.23-2.25 μmol, 25 °C, 30 min). The resulting protein extract was first diluted to 2M urea with 200 mM ammonium bicarbonate for digestion with endoproteinase LysC (1:10 w:w, 37°C, o/n, Wako, cat # 129-02541), and then diluted 2-fold with 200 mM ammonium bicarbonate for trypsin digestion (1:10 w:w, 37°C, 8h, Promega cat # V5113).

After digestion, the peptide mixes were acidified with trifluoroacetic acid (TFA) to have a final concentration of 1% TFA, desalted with a MicroSpin C18 column (The Nest Group, Inc) and dried by vacuum centrifugation. Peptide mixes were also solubilized with 100mM TEAB at a concentration of 1 μg/μl.

Samples were labeled to generate three different TMT mixes; in each TMT mix, one sample of each species was included with a common pool consisting of a mix of all the samples. Labels were used in a way that, in each TMT mix, each species uses a different TMT reagent (see Supplementary Table 8). TMT-6 label reagents were resuspended adding 41 μl of anhydrous acetonitrile (ACN) to each tube. 2 μL of the resuspended TMT label reagent was added to 10 μg of each sample. Additionally, 10 μL of anhydrous ACN and 8 μL of TEAB 100mM were added to achieve approximately a final concentration of 30% of acetonitrile. Tubes were incubated at room temperature for 1h, and then 2 μL of 5% hydroxylamine was added and incubated to quench the reaction (15 min, 25°C). Samples were combined in three mixes as shown in Supplementary Table 8, desalted with a MicroSpin C18 column (The Nest Group, Inc), and dried by vacuum centrifugation.

TMT mixes were fractionated using basic pH reversed-phase fractionation ^82^ in an Agilent 1200 system. Peptides were separated in a 90 min linear gradient from 5% to 90% acetonitrile in 10 mM ammonium bicarbonate pH 8 at 0.8 mL/min flow rate over an Agilent Zorbax Extend-C18 (4.6 mm ID, 220 mm in length, 5-m particles, 80 Å pore size). Fractions were collected every minute into a total of 96 fractions which were consolidated into 24, of which only 12 nonadjacent samples were analyzed. Fractions volume was reduced to 300 μL approximately and acidified by adding formic acid up to 10% of the final volume. Fractions were desalted with a MicroSpin C18 column (The Nest Group, Inc) and dried by vacuum centrifugation.

### Chromatographic and mass spectrometric analysis

Samples were analyzed using an LTQ-Orbitrap Fusion Lumos mass spectrometer (Thermo Fisher Scientific, San Jose, CA, USA) coupled to an EASY-nLC 1000 (Thermo Fisher Scientific (Proxeon), Odense, Denmark). Peptides were loaded directly onto the analytical column and were separated by reversed-phase chromatography using a 50-cm column with an inner diameter of 75 μm, packed with 2 μm C18 particles spectrometer (Thermo Scientific, San Jose, CA, USA).

Chromatographic gradients started at 95% buffer A and 5% buffer B with a flow rate of 300 nl/min for 5 minutes and gradually increased to 22% buffer B and 78% A in 79 min and then to 35% buffer B and 65% A in 11 min. After each analysis, the column was washed for 10 min with 10% buffer A and 90% buffer B. Buffer A: 0.1% formic acid in water. Buffer B: 0.1% formic acid in acetonitrile.

The mass spectrometer was operated in positive ionization mode with nano spray voltage set at 2.4 kV and source temperature at 275°C. Ultramark 1621 was used for external calibration of the FT mass analyzer prior to the analyses, and an internal calibration was performed using the background polysiloxane ion signal at m/z 445.12003. The dynamics exclusion duration was set at 60s, with a range in mass tolerance of ±10 ppm. Each analysis used the multinotch MS3-based TMT method ^83^. The scan sequence began with an MS1 spectrum (Orbitrap analysis; resolution 120 000; mass range 375–1500 m/z; automatic gain control (AGC) target 4 × 10^5^, maximum injection time 50 ms). In each cycle of data-dependent acquisition analysis, the most intense ions above a threshold ion count of 5000 were selected for fragmentation after each survey scan. The number of selected precursor ions for fragmentation was determined by the “Top Speed” acquisition algorithm with a cycle time set at 3s. Fragment ion spectra were produced via collision-induced dissociation (CID) at a normalized collision energy of 35% and they were acquired in the ion trap mass analyzer in “Turbo” mode. AGC was set to 10^4^, and an isolation window of 0.7 m/z and a maximum injection time of 50 ms were used. Following the acquisition of each MS2 spectrum, MS3 spectra were collected, in which multiple MS2 fragment ions are captured in the MS3 precursor population using isolation waveforms with multiple frequency notches. MS3 precursors were fragmented by high energy collision-induced dissociation (HCD) at a normalized collision energy of 65% and acquired in the Orbitrap analyzer at 50000 resolution. AGC was set to 10^5^, and an isolation window of 2 m/z and a maximum injection time of 105 ms were used. All data were acquired with Xcalibur software v4.1.31.9.

Digested bovine serum albumin (New England Biolabs cat # P8108S) was analyzed between each sample to avoid sample carryover and to assure stability of the instrument, and QCloud ^84^ has been used to control instrument longitudinal performance during the project.

### Identification of novel peptides by mass spectrometry proteomics

Acquired spectra were analyzed using the Proteome Discoverer software suite (v2.3, Thermo Fisher Scientific) and the Mascot search engine (v2.6, Matrix Science ^85^). A UniProt-based database comprising all reference proteomes from the five primates was used to identify detectable genes and narrow down the search space. After removing transcripts classified as potential artifacts by SQANTI, Iso-seq protein predictions for these genes in all species were incorporated into a customized database, together with the predicted proteins from intergenic and fusion loci. For this, NHP Iso-seq isoforms were first lifted to the most recent assembly of the corresponding species ^86^. All created target databases included a list of common contaminants ^87^ and all the corresponding decoy entries. This strategy leverages the high-quality Iso-seq isoforms detected in all species to build a comprehensive target database, which was used to identify peptides present across all samples.

For peptide identification, a precursor ion mass tolerance of 7 ppm was used for MS1 level, trypsin was chosen as the enzyme, and up to three missed cleavages were allowed. The fragment ion mass tolerance was set to 0.5 Da for MS2 spectra. Oxidation of methionine and N-terminal protein acetylation were used as variable modifications, whereas carbamidomethylation on cysteines, TMT6plex in Lysines and TMT6plex in peptide N-terminal were set as a fixed modification. The false discovery rate (FDR) in peptide identification was set to a maximum of 5%. Peptides were quantified using the reporter ion intensities in MS3 from the common peptides among the five species that were unique to a protein group. According to manufacturer’s specifications, reporter ion intensities were adjusted to correct for the isotopic impurities of the different TMT reagents. Reporter ion intensities from the samples were referenced to the common pool present in each of the three TMT mixes and they were used to estimate the peptide and protein fold-changes. The obtained fold-changes were then log-transformed and normalized between the channels by adjusting the mean log-fold-change to zero.

We established further stringent criteria to filter for high-quality peptide identifications and ensure a good quantification signal: FDR<5%, Mascot IonScore > 20, reporter ion intensity signal (abundance) per sample > 50 in any sample of a given species, and species-wise median ratio of sample abundance to pool abundance > 0.6, after discarding contaminant matches and peptides arising from more than 1 tryptic miscleavage). The detected peptides in each species were searched in the set of peptides derived from the *in-silico* digestion (tryptic digestion allowing a maximum of 1 miscleavage) of UniProt reference proteomes (based on hg38, panTro5, gorGor4, ponAbe2, and rheMac8) and RefSeq CDS set (based on hg38, panTro6, gorGor6, ponAbe3 and rheMac10) for each of the species independently. Only detected peptides present in the *in-silico* digestion of the customized database in the context of each species genome (and not in the digestions of UniProt/RefSeq proteomes) were defined as novel. Protein predictions from Iso-seq transcript models were performed using GeneMarkS-T (GMST) ^88^. Data processing was performed using EMBOSS ^89^, SeqKit ^90^, and in-house scripts. Tryptic peptides were mapped to their genomic coordinates using PGx ^91^.

### Definition of orthology relationships

Peptide sequences, gene annotations and genome assemblies were downloaded from Ensembl. Considering that one gene may contain multiple protein isoforms, the longest with a complete ORF was kept. Then, Blastp was used for peptide sequence comparison between NHP and humans with an e-value limit of 1e-5. Only reciprocal best hits (RBH) following synteny conservation were defined as orthologous. Then, we crossed our gene orthologies with Ensembl Compara one-to-one gene orthologies (V91) and kept the intersection for further analyses ^92^.

For these one-to-one orthologous genes across the five species, UCSC liftOver ^86^ was performed to identify the corresponding orthologous transcripts using available whole-genome alignments for hg38, panTro6, gorGor6, ponAbe3 and rheMac10. NHP transcript structures were projected into the novel assemblies and then into hg38 to complement human Iso-seq transcriptomes. To deal with the unbalanced isoform repertoire of Iso-seq transcriptomes (Supplementary Table 1) resulting from the lack of experimental saturation, hereafter, we will use the orthologous gene models for all species based on projected orthologous transcripts (as explained below the expression of a given transcript/exon will be assessed in every species by the corresponding species-specific RNA-seq data). The variability in internal exon ends and transcript extremes (up to 100 nucleotides) from each transcript model was collapsed using TAMA ^93^. The resulting transcript models were projected to all NHP, and we kept the models present in the five species for further analyses (species-specific gains and losses and DIU).

We followed an analogous strategy without any transcript collapsing to account for smaller splicing changes (*e.g.,* alternative donor and acceptor sites) that were quantified by DEU analyses.

### Species-specific gain and loss of exons (genomic structure)

We restricted these analyses to the set of transcripts derived from one-to-one orthologous genes. Non-human primate exons which failed to be projected into human assembly (hg38) and human exons which failed to be projected to all four NHP species by UCSC liftOver were chosen as species-specific exon candidates. Then, Liftoff ^94^ was used to perform a local alignment of the exon candidates to the other four assemblies. Only exons with an alignment rate lower than 50% in the other four assemblies were selected as species-specific exonic structures.

### Transcript expression calculation and reconstruction of transcript gains and losses

Considering the isoform discovery rate (saturation curve) did not reach the *plateau* for Iso-seq data, we incorporated deep RNA-seq (3 LCLs from each species; Human: GM12878, GM19150, GM19238; Chimpanzee: CH114, CH391, CH170; Gorilla: DIAN, OMOYE, GG05; Orangutan: PPY6 (also named PPY6_1), EB185, CRL-1850; Macaque: R02027, R05040 and R94011) to quantify the projected transcript models in each genome using Kallisto ^78^. For each sample, transcripts that were unsupported by RNA-seq in their splice junctions were assigned 0 TPM (thus excluding mono-exonic transcripts).

Count ^30^ was used to reconstruct the transcript expression gains and losses in primates considering the known phylogenetic structure and the resulting transcript expression matrix. Wagner parsimony was used with a relative penalty of gain-to-loss equal to 1. We also retrieved the number of transcript gains and losses that can be explained by a unique gain or loss event in our phylogeny and compared them to the corresponding gains and losses inferred by Wagner parsimony reconstruction for the same branches by calculating the ratio between them.

To account for the possible effect of intra-species transcript expression variability in the detection of transcript gains and losses, we further asked for strict intra-species consistency in the presence/absence of transcript expression. Hence, we required that the 3 biological replicates (different LCLs) from each species are coherent in the expression/absence pattern of each transcript. Thus, species-specific transcripts (isoform innovations) are expressed in all LCLs from a given species and absent in the rest of LCLs from the remaining species.

The proportion of different alternative splicing classes in species-specific transcripts was compared to that in conserved transcripts (SUPPA definition, see ‘Classification of alternative splicing patterns’). We excluded RI from this comparison since species-specific RI events are less detectable considering our quantification strategy relies on RNA-seq support in transcript splice junctions, without considering the RNA-seq coverage in the boundary between an exon and a retained intron.

### Differential gene expression analyses

To assess the effect of gene expression in the detection of species-specific transcripts, differential gene expression analyses were performed using DESeq2 (pairwise comparatives) ^95^, including the 3 RNA-seq biological replicates per species. Genes with 10 or more RNA-seq reads accumulated across samples were retained. Species-specific upregulated genes were defined as those showing significant overexpression (permissive padj<0.1 and log2FoldChange>0 were used to detect even subtle influences of gene expression changes in isoform detection) in a given species compared to all the rest, regardless of the possible gene expression differences among the remaining species. Then, genes upregulated in each species were intersected with those expressing species-specific transcripts in the corresponding species.

We followed an analogous strategy to define species-specific downregulated genes (padj<0.1, log2FoldChange<0). Species-specific up and downregulated genes were intersected with those displaying species-specific up or downregulated exon usage in the same species.

### Differential isoform usage analyses

Kallisto transcript quantifications were used to evaluate interspecies isoform usage (IU) changes ^96^. ComBat batch effect correction ^97^ and TMM (Trimmed Mean of M-values) normalization were performed prior to this calculation. IU values for orthologous transcripts were calculated across all samples using IsoformSwitchAnalyzeR package ^98^. Then, principal component analysis (PCA) and hierarchical clustering based on the euclidean distances of Spearman correlations for IU across samples were computed.

To measure the most confident isoform usage changes, we first excluded genes in which the transcript models resulted from the collapsing of internal exon boundaries, since this variability in internal exon ends can lead to artificial isoform usage differences considering our quantification strategy (see ‘Transcript expression calculation and reconstruction of transcript gains and losses’). In addition, low expression genes/isoforms and genes with a single isoform were removed using IsoformSwitchAnalyzeR preFilter function (default settings: geneExpressionCutoff = 1 TPM, isoformExpressionCutoff = 0 TPM, IFcutoff = 0.01, removeSingleIsoformGenes = TRUE). Differential isoform usage (DIU) analyses were performed by establishing pairwise comparatives across the five species using DEXSeq method in IsoformSwitchAnalyzeR ^99–101^. Species-specific DIU cases were defined as significant isoform usage changes (|dIU|>0.1 and isoform switch q-value<0.05) in a given species compared to the rest of primates (up or downregulated), while the remaining species showed non-significant differences among them. The same rationale was applied to classify isoform usage changes shared by groups of 2 or 3 species in comparison to the rest of them. IsoformSwitchAnalyzeR preFilter function (default settings) was also used for the evaluation of genes where orthologous splice junctions showed high-impact genetic changes across primates.

### Differential exon usage analyses

The projected transcript models in each species (without any collapsing, see ‘Definition of orthology relationships’) were flattened to define orthologous exonic counting bins from all transcribed segments (*i.e.*, exonic parts). RNA-seq read counts overlapping each exonic part in the 3 LCLs from each species (biological replicates) were obtained using HTSeq-count ^102^. Differential exon usage (DEU) analyses were performed by establishing pairwise comparatives across the five species using DEXSeq ^101,103^ while controlling for batch effects. Significant exon usage changes (exonBaseMean>10, |log2FC|>1.2 and padj<0.05) were retrieved from each pairwise comparative and used to classify the change pattern across species. Exonic parts showing significant difference in usage in a given species compared to the rest of primates (up or downregulated) were defined as species-specific as long as the remaining species did not display significant differences among them. The same rationale was used to classify changes shared by groups of 2 or 3 species compared to the remaining species. Exonic parts not classified as DEU in any pairwise comparative and displaying exonBaseMean>10 (average across all samples) were defined as conserved.

To quantify the frequency of exon inclusion, we calculated percent-spliced-in (PSI) estimates according to ^104^ and computed the average across the 15 RNA-seq experiments (3 LCLs per species). Average PSI estimates were used to classify exonic parts into highly excluded [0-0.2), middle excluded [0.2-0.4), middle included [0.4-0.8) and highly included [0.8-1] (Fig. 5a). We classified the exonic parts into AS modes using SUPPA in the projected transcript models (see ‘Classification of alternative splicing patterns’). The phastCons conservation scores across primates were retrieved from UCSC whole-genome alignments (17-way phastCons scores). Average phastCons scores per exonic part were classified into low [0-0.3], middle (0.3-0.8) and high [0.8-1] (Fig. 5a). Nucleotide diversity (pi) in human populations was calculated from the data generated by the 1000 Genomes project ^105^ mapped against hg38 using VCFtools ^106^. The 1000 Genomes strict mask was used to select exonic parts with all their nucleotides classified as passed bases according to the strict mask BED file (hg38). Average pi estimates were also computed for each exonic part.

### Classification of genes according to their splicing and usage patterns

One-to-one orthologous genes expressed in LCLs were classified according to the splicing and usage changes detected in the analyses of transcript expression, DIU, and DEU.

First of all, isoforms and exonic parts were classified as conserved (if the expression or usage is conserved in all species), species-specific gains (upregulation in a single-species in comparison to the rest), and other changes (*i.e.*, the remaining cases resulting from expression or usage differences between groups of species, or species-specific losses). Isoforms and exonic parts showing usage patterns that are not consistent with differences between groups of species (*e.g.,* differential usage in only one pairwise comparative) were not considered as they are more likely to reflect intra-species variability.

The classes of isoforms and exonic parts resulting from the analyses of transcript expression, DIU, and DEU were aggregated at the gene level. Each gene was classified according to the following rules:

1. ‘all_conserved’: genes where only conserved isoforms/exonic parts were detected.
2. ‘only_human_sp’: genes where human-specific gains were detected (and not species-specific gains from other species), allowing the presence of conserved or other isoforms/exonic parts.
3. ‘only_NHP_sp’: genes where only chimpanzee, gorilla, orangutan or macaque-specific gains were detected (for a single species, excluding human), allowing the presence of conserved or other isoforms/exonic parts.
4. ‘convergence_spsp’: genes where independent events of species-specific gains were detected in multiple species, allowing the presence of conserved or other isoforms/exonic parts.
5. ‘other’: genes where other patterns were detected (*i.e.*, not species-specific gains), allowing the presence of conserved isoforms/exonic parts.

### dN/dS in human populations and positive selection in primates

Pre-computed ratios of nonsynonymous to synonymous substitutions (dN/dS) in human populations and genes undergoing positive selection in the primate lineage were retrieved from ^55^. Statistical significance for the dN/dS differences (pairwise comparisons across gene classes) was obtained by Dwass-Steel-Critchlow-Fligner all-pairs test, with a single-step p-value adjustment. Only genes with precomputed dN/dS were included in this comparative. Enrichment in positively selected genes across gene classes (in comparison to genes with fully conserved splicing and usage patterns) was assessed using Fisher’s exact test (Bonferroni p-value adjustment) after restricting to the background genes defined in ^55^.

### Functional enrichment analysis

Functional enrichment was performed using over-representation analysis (ORA) in WebGestalt ^107^ based on Gene Ontology Biological Processes ^108,109^ and Panther pathways ^110^ . We adjusted the background gene set to the orthologous genes expressed in LCLs under evaluation in each analysis for each functional enrichment. In addition, tissue-specific gene expression was tested using tissueEnrich ^111^.

We tested the enrichment and depletion in particular gene lists related to the immune response, immune disease, LCL-specific, and housekeeping genes (Fisher’s exact test). Immune response genes belong to GO:0006955 and their child terms and were obtained from Ensembl Biomart V91. A manually curated classification of innate immunity genes (8 subgroups: accessory molecule, adaptor, effector, regulator, secondary receptor, sensor, signal transducer, and transcription genes) was retrieved from ^31^. Immune disease genes correspond to those provided by the International Union of Immunological Societies (IUIS, December 2020) ^112^. Genes with LCL-specific expression patterns were downloaded from ^113^, and housekeeping genes were retrieved from the Housekeeping and Reference Transcript Atlas ^114^.

## Acknowledgements

This work was supported by the Strategic Priority Research Program of the Chinese Academy of Sciences (XDB31020000); the National Key R&D Program of China (MOST) grant 2018YFC1406901; the International Partnership Program of the Chinese Academy of Sciences (no. 152453KYSB20170002); the Carlsberg Foundation (CF16-0663); the Villum Foundation (no. 25900) to G.Z.; La Caixa Foundation (ID 100010434) Fellowship Code LCF/BQ/DE16/11570011 (L.F.-P.). The CRG/UPF Proteomics Unit is part of the Spanish Infrastructure for Omics Technologies (ICTS OmicsTech) and it is member of ProteoRed PRB3 consortium which is supported by grant PT17/0019 of the PE I+D+i 2013-2016 from the Instituto de Salud Carlos III (ISCIII), ERDF, and “Secretaria d’Universitats i Recerca del Departament d’Economia i Coneixement de la Generalitat de Catalunya” (2017SGR595). We thank China National GeneBank for providing the computational resources. TMB is supported by funding from the European Research Council (ERC) under the European Union’s Horizon 2020 research and innovation programme (grant agreement No. 864203), BFU2017-86471-P (MINECO/FEDER, UE), “Unidad de Excelencia María de Maeztu”, funded by the AEI (CEX2018-000792-M), Howard Hughes International Early Career, NIH 1R01HG010898-01A1 and Secretaria d’Universitats i Recerca and CERCA Programme del Departament d’Economia i Coneixement de la Generalitat de Catalunya (GRC 2017 SGR 880).

## Ethics declarations

### Availability of data and materials

The datasets supporting the conclusions of this article will be available in the SRA and PRIDE repositories.

## References

1. Bush, S. J., Chen, L., Tovar-Corona, J. M. & Urrutia, A. O. Alternative splicing and the evolution of phenotypic novelty. Philos. Trans. R. Soc. Lond. B Biol. Sci. 372, 20150474 (2017).

2. Scotti, M. M. & Swanson, M. S. RNA mis-splicing in disease. Nature Reviews Genetics 17, 19–32 (2016).

3. Zhang, S.-J. et al. Isoform Evolution in Primates through Independent Combination of Alternative RNA Processing Events. Mol. Biol. Evol. 34, 2453–2468 (2017).

4. Dougherty, M. L. et al. Transcriptional fates of human-specific segmental duplications in brain. Genome Res. 28, 1566–1576 (2018).

5. Kronenberg, Z. N. et al. High-resolution comparative analysis of great ape genomes. Science 360, eaar6343 (2018).

6. Xiong, J. et al. Predominant patterns of splicing evolution on human, chimpanzee and macaque evolutionary lineages. Hum. Mol. Genet. 27, 1474–1485 (2018).

7. Mazin, P. V. et al. Conservation, evolution, and regulation of splicing during prefrontal cortex development in humans, chimpanzees, and macaques. RNA 24, 585–596 (2018).

8. Mazin, P. et al. Widespread splicing changes in human brain development and aging. Mol. Syst. Biol. 9, 633 (2013).

9. Blekhman, R., Marioni, J. C., Zumbo, P., Stephens, M. & Gilad, Y. Sex-specific and lineage-specific alternative splicing in primates. Genome Res. 20, 180–189 (2010).

10. Mittleman, B. E. et al. Divergence in alternative polyadenylation contributes to gene regulatory differences between humans and chimpanzees. Elife 10, e62548 (2021).

11. Agosto, L. M. et al. Deep profiling and custom databases improve detection of proteoforms generated by alternative splicing. Genome Res. 29, 2046–2055 (2019).

12. Liu, Y. et al. Impact of Alternative Splicing on the Human Proteome. Cell Rep. 20, 1229–1241 (2017).

13. Sheynkman, G. M., Shortreed, M. R., Frey, B. L. & Smith, L. M. Discovery and Mass Spectrometric Analysis of Novel Splice-junction Peptides Using RNA-Seq. Molecular & Cellular Proteomics 12, 2341–2353 (2013).

14. Tress, M. L., Abascal, F. & Valencia, A. Alternative Splicing May Not Be the Key to Proteome Complexity. Trends in Biochemical Sciences 42, 98–110 (2017).

15. Abascal, F. et al. Alternatively Spliced Homologous Exons Have Ancient Origins and Are Highly Expressed at the Protein Level. PLoS Comput. Biol. 11, e1004325 (2015).

16. Carlyle, B. C. et al. Isoform-Level Interpretation of High-Throughput Proteomics Data Enabled by Deep Integration with RNA-seq. J. Proteome Res. 17, 3431–3444 (2018).

17. Lau, E. et al. Splice-Junction-Based Mapping of Alternative Isoforms in the Human Proteome. Cell Rep. 29, 3751–3765.e5 (2019).

18. Mignone, F., Gissi, C., Liuni, S. & Pesole, G. Untranslated regions of mRNAs. Genome Biol. 3, 1–10 (2002).

19. Byrne, A., Cole, C., Volden, R. & Vollmers, C. Realizing the potential of full-length transcriptome sequencing. Philos. Trans. R. Soc. Lond. B Biol. Sci. 374, 20190097 (2019).

20. He, Y. et al. Long-read assembly of the Chinese rhesus macaque genome and identification of ape-specific structural variants. Nat. Commun. 10, 4233 (2019).

21. Warren, W. C. et al. Sequence diversity analyses of an improved rhesus macaque genome enhance its biomedical utility. Science 370, eabc6617 (2020).

22. Mao, Y. et al. A high-quality bonobo genome refines the analysis of hominid evolution. Nature 594, 77–81 (2021).

23. Tardaguila, M. et al. SQANTI: extensive characterization of long-read transcript sequences for quality control in full-length transcriptome identification and quantification. Genome Res. 28, 396–411 (2018).

24. Bundred, J. R., Hendrix, E. & Coleman, M. L. The emerging roles of ribosomal histidyl hydroxylases in cell biology, physiology and disease. Cell. Mol. Life Sci. 75, 4093–4105 (2018).

25. Tokita, M. J. et al. De Novo Truncating Variants in SON Cause Intellectual Disability, Congenital Malformations, and Failure to Thrive. The American Journal of Human Genetics 99, 720–727 (2016).

26. Kim, J.-H. et al. De Novo Mutations in SON Disrupt RNA Splicing of Genes Essential for Brain Development and Metabolism, Causing an Intellectual-Disability Syndrome. Am. J. Hum. Genet. 99, 711–719 (2016).

27. Sun, C. T. et al. Transcription repression of human hepatitis B virus genes by negative regulatory element-binding protein/SON. J. Biol. Chem. 276, 24059–24067 (2001).

28. Ahn, E.-Y. et al. SON controls cell-cycle progression by coordinated regulation of RNA splicing. Mol. Cell 42, 185–198 (2011).

29. Cordaux, R. & Batzer, M. A. The impact of retrotransposons on human genome evolution. Nature Reviews Genetics 10, 691–703 (2009).

30. Csűös, M. Count: evolutionary analysis of phylogenetic profiles with parsimony and likelihood. Bioinformatics 26, 1910–1912 (2010).

31. Deschamps, M. et al. Genomic Signatures of Selective Pressures and Introgression from Archaic Hominins at Human Innate Immunity Genes. Am. J. Hum. Genet. 98, 5–21 (2016).

32. Mathis, B. J., Lai, Y., Qu, C., Janicki, J. S. & Cui, T. CYLD-mediated signaling and diseases. Curr. Drug Targets 16, 284–294 (2015).

33. Sun, S.-C. CYLD: a tumor suppressor deubiquitinase regulating NF-κB activation and diverse biological processes. Cell Death & Differentiation 17, 25–34 (2010).

34. Srokowski, C. C. et al. Naturally occurring short splice variant of CYLD positively regulates dendritic cell function. Blood 113, 5891–5895 (2009).

35. Hövelmeyer, N. et al. Regulation of B cell homeostasis and activation by the tumor suppressor gene CYLD. J. Exp. Med. 204, 2615–2627 (2007).

36. Shaath, H., Vishnubalaji, R., Elkord, E. & Alajez, N. M. Single-Cell Transcriptome Analysis Highlights a Role for Neutrophils and Inflammatory Macrophages in the Pathogenesis of Severe COVID-19. Cells 9, 2374 (2020).

37. Herberg, J. A. et al. Diagnostic Test Accuracy of a 2-Transcript Host RNA Signature for Discriminating Bacterial vs Viral Infection in Febrile Children. JAMA 316, 835–845 (2016).

38. Busse, D. C. et al. Interferon-Induced Protein 44 and Interferon-Induced Protein 44-Like Restrict Replication of Respiratory Syncytial Virus. J. Virol. 94, e00297–20 (2020).

39. Li, Y. et al. IFI44L expression is regulated by IRF-1 and HIV-1. FEBS Open Bio 11, 105–113 (2020).

40. Bian, P. et al. RIPK3 Promotes JEV Replication in Neurons Downregulation of IFI44L. Front. Microbiol. 11, 368 (2020).

41. Hadjadj, J. et al. Impaired type I interferon activity and inflammatory responses in severe COVID-19 patients. Science 369, 718–724 (2020).

42. Mick, E. et al. Upper airway gene expression differentiates COVID-19 from other acute respiratory illnesses and reveals suppression of innate immune responses by SARS-CoV-2. medRxiv (2020) doi:10.1101/2020.05.18.20105171.

43. de Oyarzabal, E. et al. Expression of USP18 and IL2RA Is Increased in Individuals Receiving Latent Tuberculosis Treatment with Isoniazid. J Immunol Res 2019, 1297131 (2019).

44. Unterman, A. et al. Single-Cell Omics Reveals Dyssynchrony of the Innate and Adaptive Immune System in Progressive COVID-19. medRxiv (2020) doi:10.1101/2020.07.16.20153437.

45. Zhao, M. et al. IFI44L promoter methylation as a blood biomarker for systemic lupus erythematosus. Ann. Rheum. Dis. 75, 1998–2006 (2016).

46. Lempainen, J. et al. Associations Between IFI44L Gene Variants and Rates of Respiratory Tract Infections During Early Childhood. J. Infect. Dis. 223, 157–165 (2021).

47. Haralambieva, I. H. et al. Genome-wide associations of CD46 and IFI44L genetic variants with neutralizing antibody response to measles vaccine. Hum. Genet. 136, 421–435 (2017).

48. Feenstra, B. et al. Common variants associated with general and MMR vaccine-related febrile seizures. Nat. Genet. 46, 1274–1282 (2014).

49. Wu, T., Ren, M.-X., Chen, G.-P., Jin, Z.-M. & Wang, G. Rrp15 affects cell cycle, proliferation, and apoptosis in NIH3T3 cells. FEBS Open Bio 6, 1085–1092 (2016).

50. Kuhlwilm, M. & Boeckx, C. A catalog of single nucleotide changes distinguishing modern humans from archaic hominins. Sci. Rep. 9, 8463 (2019).

51. Zhao, B. et al. DescribePROT: database of amino acid-level protein structure and function predictions. Nucleic Acids Res. 49, D298–D308 (2020).

52. Zhang, J. & Kurgan, L. SCRIBER: accurate and partner type-specific prediction of protein-binding residues from proteins sequences. Bioinformatics 35, i343–i353 (2019).

53. Benjamini, Y., Drai, D., Elmer, G., Kafkafi, N. & Golani, I. Controlling the false discovery rate in behavior genetics research. Behav. Brain Res. 125, 279–284 (2001).

54. Soneson, C., Matthes, K. L., Nowicka, M., Law, C. W. & Robinson, M. D. Isoform prefiltering improves performance of count-based methods for analysis of differential transcript usage. Genome Biol. 17, 12 (2016).

55. van der Lee, R., Wiel, L., van Dam, T. J. P. & Huynen, M. A. Genome-scale detection of positive selection in nine primates predicts human-virus evolutionary conflicts. Nucleic Acids Res. 45, 10634–10648 (2017).

56. Nelms, K., Keegan, A. D., Zamorano, J., Ryan, J. J. & Paul, W. E. The IL-4 receptor: signaling mechanisms and biologic functions. Annu. Rev. Immunol. 17, 701–738 (1999).

57. Payer, L. M. et al. Alu insertion variants alter mRNA splicing. Nucleic Acids Res. 47, 421–431 (2019).

58. Song, G. et al. New centromere autoantigens identified in systemic sclerosis using centromere protein microarrays. J. Rheumatol. 40, 461–468 (2013).

59. Bancroft, J., Auckland, P., Samora, C. P. & McAinsh, A. D. Chromosome congression is promoted by CENP-Q- and CENP-E-dependent pathways. J. Cell Sci. 128, 171–184 (2015).

60. Rao, S. S. P. et al. A 3D Map of the Human Genome at Kilobase Resolution Reveals Principles of Chromatin Looping. Cell 162, 687–688 (2015).

61. Moreno-De-Luca, D. et al. Deletion 17q12 Is a Recurrent Copy Number Variant that Confers High Risk of Autism and Schizophrenia. Am. J. Hum. Genet. 87, 618–630 (2010).

62. Nagamani, S. C. S. et al. Clinical spectrum associated with recurrent genomic rearrangements in chromosome 17q12. Eur. J. Hum. Genet. 18, 278–284 (2010).

63. Mefford, H. C. et al. Recurrent reciprocal genomic rearrangements of 17q12 are associated with renal disease, diabetes, and epilepsy. Am. J. Hum. Genet. 81, 1057–1069 (2007).

64. Cardone, M. F. et al. Hominoid chromosomal rearrangements on 17q map to complex regions of segmental duplication. Genome Biol. 9, R28 (2008).

65. Rodriguez, J. M. et al. APPRIS 2017: principal isoforms for multiple gene sets. Nucleic Acids Res. 46, D213–D217 (2018).

66. Ezkurdia, I. et al. Most highly expressed protein-coding genes have a single dominant isoform. J. Proteome Res. 14, 1880–1887 (2015).

67. Barreiro, L. B. & Quintana-Murci, L. From evolutionary genetics to human immunology: how selection shapes host defence genes. Nat. Rev. Genet. 11, 17–30 (2010).

68. Barreiro, L. B., Marioni, J. C., Blekhman, R., Stephens, M. & Gilad, Y. Functional comparison of innate immune signaling pathways in primates. PLoS Genet. 6, e1001249 (2010).

69. Sams, A. J. et al. Adaptively introgressed Neandertal haplotype at the OAS locus functionally impacts innate immune responses in humans. Genome Biol. 17, 246 (2016).

70. Rotival, M., Quach, H. & Quintana-Murci, L. Defining the genetic and evolutionary architecture of alternative splicing in response to infection. Nat. Commun. 10, 1671 (2019).

71. Mayr, C. What Are 3’ UTRs Doing? Cold Spring Harb. Perspect. Biol. 11, 10 a034728 (2019).

72. García-Pérez, R. et al. Epigenomic profiling of primate lymphoblastoid cell lines reveals the evolutionary patterns of epigenetic activities in gene regulatory architectures. Nat. Commun. 12, 1–17 (2021).

73. Chen, Y. et al. SOAPnuke: a MapReduce acceleration-supported software for integrated quality control and preprocessing of high-throughput sequencing data. Gigascience 7, 1–6 (2018).

74. Dobin, A. et al. STAR: ultrafast universal RNA-seq aligner. Bioinformatics 29, 15–21 (2013).

75. Salmela, L. & Rivals, E. LoRDEC: accurate and efficient long read error correction. Bioinformatics 30, 3506–3514 (2014).

76. Wu, T. D. & Watanabe, C. K. GMAP: a genomic mapping and alignment program for mRNA and EST sequences. Bioinformatics 21, 1859–1875 (2005).

77. Wyman, D. & Mortazavi, A. TranscriptClean: variant-aware correction of indels, mismatches and splice junctions in long-read transcripts. Bioinformatics 35, 340–342 (2019).

78. Bray, N. L., Pimentel, H., Melsted, P. & Pachter, L. Near-optimal probabilistic RNA-seq quantification. Nat. Biotechnol. 34, 525–527 (2016).

79. Li, H. et al. The Sequence Alignment/Map format and SAMtools. Bioinformatics 25, 2078–2079 (2009).

80. Quinlan, A. R. & Hall, I. M. BEDTools: a flexible suite of utilities for comparing genomic features. Bioinformatics 26, 841–842 (2010).

81. Alamancos, G. P., Pagès, A., Trincado, J. L., Bellora, N. & Eyras, E. Leveraging transcript quantification for fast computation of alternative splicing profiles. RNA 21, 1521–1531 (2015).

82. Isasa, M. et al. Multiplexed, Proteome-Wide Protein Expression Profiling: Yeast Deubiquitylating Enzyme Knockout Strains. J. Proteome Res. 14, 5306–5317 (2015).

83. McAlister, G. C. et al. MultiNotch MS3 enables accurate, sensitive, and multiplexed detection of differential expression across cancer cell line proteomes. Anal. Chem. 86, 7150–7158 (2014).

84. Chiva, C. et al. QCloud: A cloud-based quality control system for mass spectrometry-based proteomics laboratories. PLoS One 13, e0189209 (2018).

85. Perkins, D. N., Pappin, D. J., Creasy, D. M. & Cottrell, J. S. Probability-based protein identification by searching sequence databases using mass spectrometry data. Electrophoresis 20, 3551–3567 (1999).

86. Hinrichs, A. S. et al. The UCSC Genome Browser Database: update 2006. Nucleic Acids Res. 34, D590–598 (2006).

87. Beer, L. A., Liu, P., Ky, B., Barnhart, K. T. & Speicher, D. W. Efficient Quantitative Comparisons of Plasma Proteomes Using Label-Free Analysis with MaxQuant. Methods Mol. Biol. 1619, 339–352 (2017).

88. Tang, S., Lomsadze, A. & Borodovsky, M. Identification of protein coding regions in RNA transcripts. Proceedings of the 5th ACM Conference on Bioinformatics, Computational Biology, and Health Informatics – BCB ’14 43, e78–e78 (2014).

89. Rice, P., Longden, I. & Bleasby, A. EMBOSS: The European Molecular Biology Open Software Suite. Trends in Genetics 16, 276–277 (2000).

90. Shen, W., Le, S., Li, Y. & Hu, F. SeqKit: A Cross-Platform and Ultrafast Toolkit for FASTA/Q File Manipulation. PLOS ONE 11, e0163962 (2016).

91. Askenazi, M., Ruggles, K. V. & Fenyö, D. PGx: Putting Peptides to BED. J. Proteome Res. 15, 795–799 (2016).

92. Yates, A. D. et al. Ensembl 2020. Nucleic Acids Res. 48, D682–D688 (2020).

93. Kuo, R. I. et al. Illuminating the dark side of the human transcriptome with long read transcript sequencing. BMC Genomics 21, 751 (2020).

94. Shumate, A. & Salzberg, S. L. Liftoff: accurate mapping of gene annotations. Bioinformatics 37, 1639–1643 (2020).

95. Love, M. I., Huber, W. & Anders, S. Moderated estimation of fold change and dispersion for RNA-seq data with DESeq2. Genome Biol. 15, 550 (2014).

96. Soneson, C., Love, M. I. & Robinson, M. D. Differential analyses for RNA-seq: transcript-level estimates improve gene-level inferences. F1000Res. 4, 1521 (2015).

97. Zhang, Y., Parmigiani, G. & Evan Johnson, W. ComBat-Seq: batch effect adjustment for RNA-Seq count data. NAR Genomics and Bioinformatics 2, 3 lqaa078 (2020).

98. Vitting-Seerup, K. & Sandelin, A. IsoformSwitchAnalyzeR: analysis of changes in genome-wide patterns of alternative splicing and its functional consequences. Bioinformatics 35, 4469–4471 (2019).

99. Vitting-Seerup, K. & Sandelin, A. The Landscape of Isoform Switches in Human Cancers. Mol. Cancer Res. 15, 1206–1220 (2017).

100. Ritchie, M. E. et al. limma powers differential expression analyses for RNA-sequencing and microarray studies. Nucleic Acids Res. 43, e47 (2015).

101. Anders, S., Reyes, A. & Huber, W. Detecting differential usage of exons from RNA-seq data. Genome Res. 22, 2008–2017 (2012).

102. Anders, S., Pyl, P. T. & Huber, W. HTSeq--a Python framework to work with high-throughput sequencing data. Bioinformatics 31, 166–169 (2015).

103. Reyes, A. et al. Drift and conservation of differential exon usage across tissues in primate species. Proc. Natl. Acad. Sci. U. S. A. 110, 15377–15382 (2013).

104. Schafer, S. et al. Alternative Splicing Signatures in RNA-seq Data: Percent Spliced in (PSI). Curr. Protoc. Hum. Genet. 87, 11.16.1–11.16.14 (2015).

105. Consortium, T. 1000 G. P. & The 1000 Genomes Project Consortium. A global reference for human genetic variation. Nature 526, 68–74 (2015).

106. Danecek, P. et al. The variant call format and VCFtools. Bioinformatics 27, 2156–2158 (2011).

107. Liao, Y., Wang, J., Jaehnig, E. J., Shi, Z. & Zhang, B. WebGestalt 2019: gene set analysis toolkit with revamped UIs and APIs. Nucleic Acids Res. 47, W199–W205 (2019).

108. Ashburner, M. et al. Gene ontology: tool for the unification of biology. The Gene Ontology Consortium. Nat. Genet. 25, 25–29 (2000).

109. Gene Ontology Consortium. The Gene Ontology resource: enriching a GOld mine. Nucleic Acids Res. 49, D325–D334 (2021).

110. Mi, H. et al. PANTHER version 16: a revised family classification, tree-based classification tool, enhancer regions and extensive API. Nucleic Acids Res. 49, D394–D403 (2020).

111. Jain, A. & Tuteja, G. TissueEnrich: Tissue-specific gene enrichment analysis. Bioinformatics 35, 1966–1967 (2019).

112. Tangye, S. G. et al. Human Inborn Errors of Immunity: 2019 Update on the Classification from the International Union of Immunological Societies Expert Committee. J. Clin. Immunol. 40, 24–64 (2020).

113. Ryaboshapkina, M. & Hammar, M. Tissue-specific genes as an underutilized resource in drug discovery. Sci. Rep. 9, 7233 (2019).

114. Hounkpe, B. W., Chenou, F., de Lima, F. & De Paula, E. V. HRT Atlas v1.0 database: redefining human and mouse housekeeping genes and candidate reference transcripts by mining massive RNA-seq datasets. Nucleic Acids Research 49, D947–D955 (2020).

