## Supplementary Information file for "Transcriptome innovations in primates revealed by single-molecule long-read sequencing"

This file contains the description of:

- **Supplementary Figures 1-20**
- **Supplementary Tables 1-8**
- **Supplementary Data 1-10**
- **Supplementary References**

### Supplementary Figures

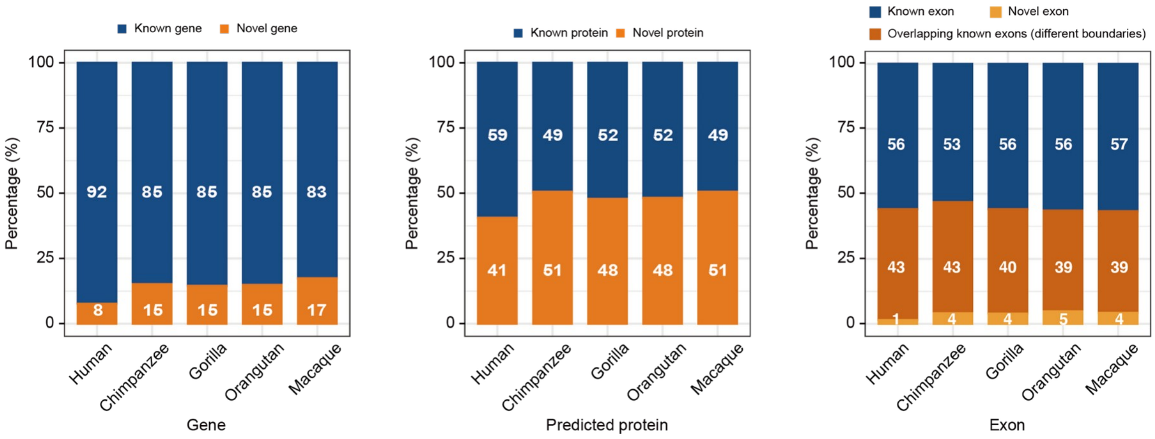


**Supplementary Figure 1.** Percentage of known vs novel genes (left), predicted proteins sequences (middle) and exons (right) captured by Iso-seq in each species.

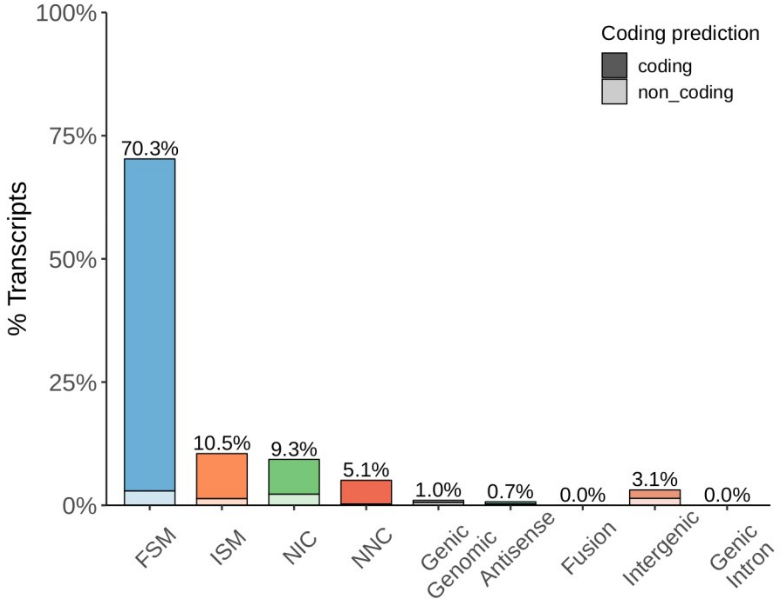


**Supplementary Figure 2.** Percentage of human Iso-seq transcripts belonging to each SQANTI structural category based on the Universal Human Reference RNA (UHRR) Iso-seq dataset. Legend indicates if the transcript has been predicted to encode a protein ('coding') or not ('non_coding'). FSM: full splice match; ISM: incomplete splice match; NIC: novel in catalog; NNC: novel not in catalog. A detailed description of SQANTI structural categories can be found in [1].

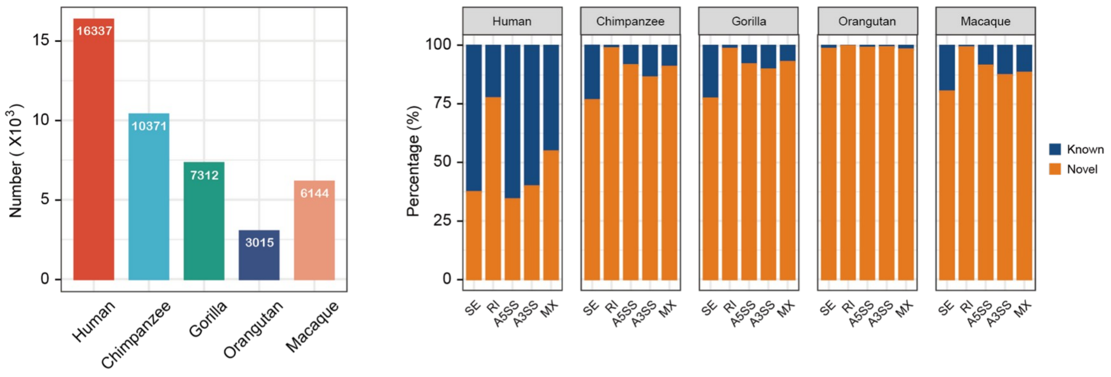


**Supplementary Figure 3.** Number of alternative splicing events (left) and percentage of known vs novel alternative splicing events captured by Iso-seq in each species (right). SE: skipping exons; RI: retained introns; A5SS: alternative 5’ splice sites; A3SS: alternative 3’ splice sites; MX: mutually exclusive exons.

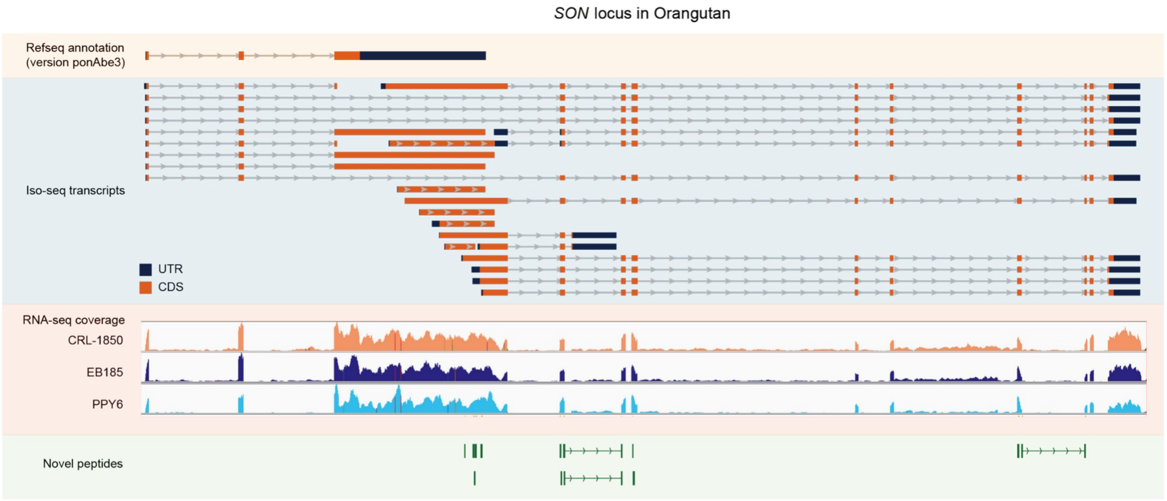


**Supplementary Figure 4.** Example of poorly annotated gene, *SON* (shown in ponAbe3 assembly), refined by Iso-seq, RNA-seq and mass spectrometry in orangutan samples.

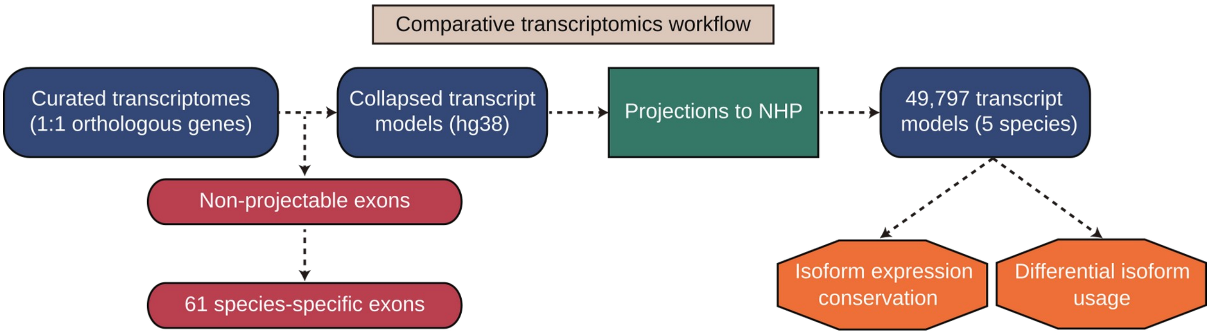


**Supplementary Figure 5.** Schematic workflow for comparative transcriptomics analyses based on Iso-seq collapsed isoform models.

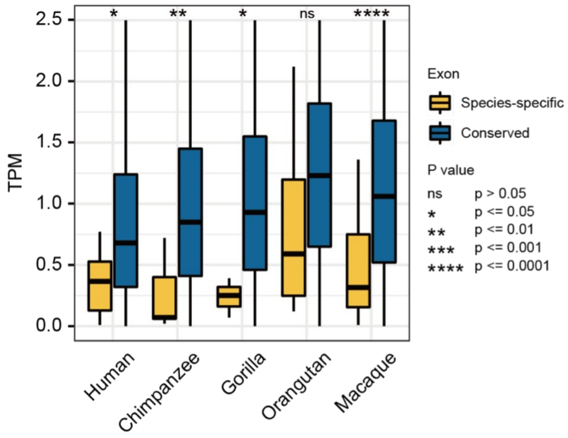


**Supplementary Figure 6.** Expression values for the exons present in a single species (Species-specific) and present across all species (Conserved). Exon expression was normalized to Transcripts Per Million (TPM). Statistical significance of the difference across groups was assessed by Student t-test.

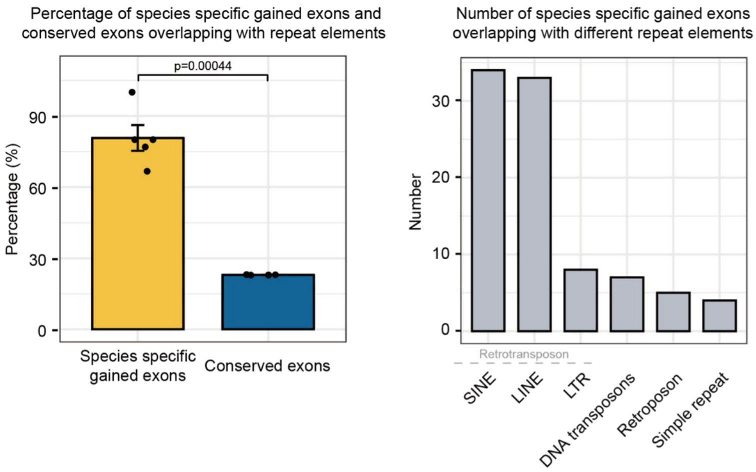


**Supplementary Figure 7.** Percentage of species-specific gained (yellow) and conserved (blue) exons overlapping repeat elements (left) and number of species-specific gained exons overlapping different classes of repeat elements (right). Statistical significance of the difference across groups was assessed by Student t-test.

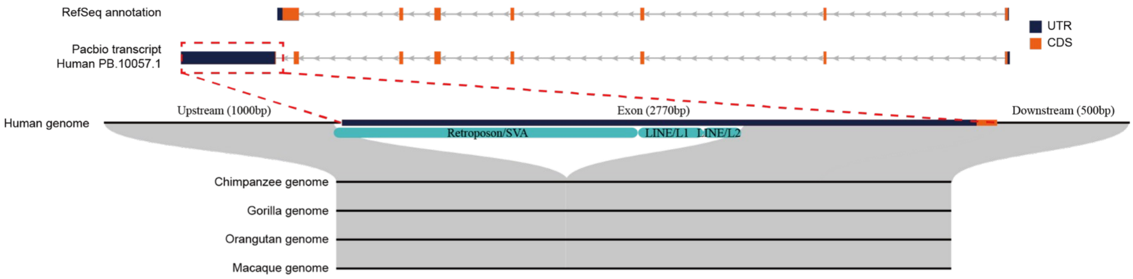


**Supplementary Figure 8.** Example of a gene, *MRNIP*, expressing a human-specific exon in 3’ UTR. RefSeq *MRNIP* models in hg38 assembly (top track) are illustrated together with human PB.10057.1 Iso-seq transcript (middle track). Human-specific insertions of transposable elements (SVA and LINEs) in 3’ UTR are also depicted (bottom track). UTR and CDS are colored in blue and orange, respectively. CDS: coding sequence; UTR: untranslated region.

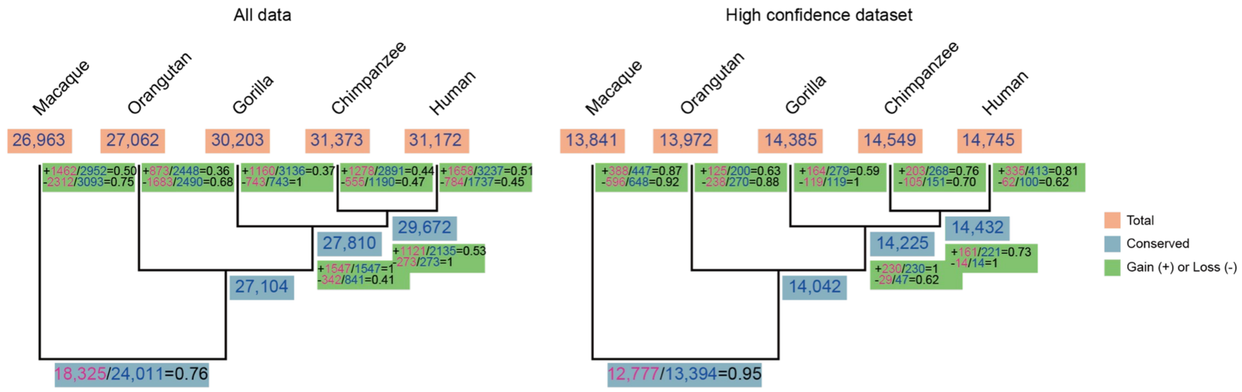


**Supplementary Figure 9.** Transcript gains and losses in the primate lineage in all projected transcripts (‘All data’, left) and high confidence transcripts (‘High confidence dataset’, corresponding to transcripts whose expression is consistent in all samples from each species, right). Numbers in blue indicate the gains and losses inferred from Wagner parsimony, whereas numbers in pink correspond to the number of transcript gains and losses that can be explained by a unique gain or loss event in our phylogeny.

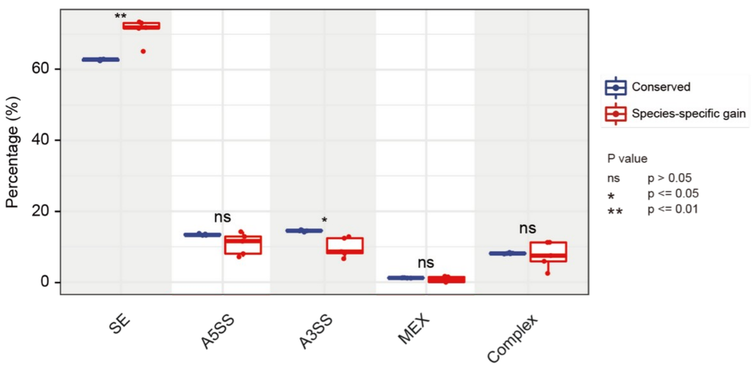


**Supplementary Figure 10.** Percentage of alternative splicing modes in transcripts expressed in all species (‘Conserved’, blue boxes) and transcripts with species-specific expression (‘Species-specific gain’, red boxes). SE, skipping exons; A5SS, alternative 5’ splice site; A3SS, alternative 3’ splice site; MEX, mutually exclusive exons; Complex, any combination of the previous. Statistical significance of the difference across groups was obtained by t.test. Retained introns (RI) are excluded from this analysis (see Methods).

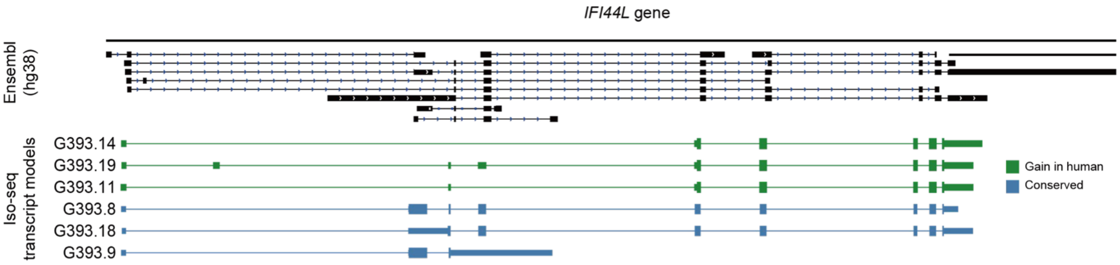


**Supplementary Figure 11.** Example of a gene, *IFI44L*, expressing human-specific and conserved transcripts. Ensembl annotation in hg38 assembly (black) is displayed (top) together with *IFI44L* Iso-seq transcript models (bottom) that are human-specific (green) or expressed in all species (blue). In Iso-seq transcript models, narrow blocks represent the untranslated region (UTR) and thick blocks indicate the coding region (CDS).

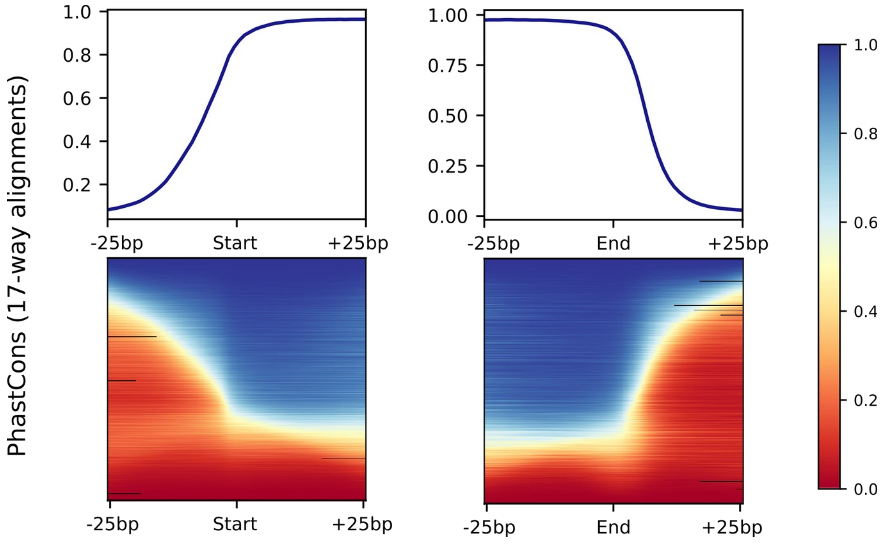


**Supplementary Figure 12.** PhastCons conservation scores in 5’ (Start) and 3’ (End) splice sites and surrounding regions (± 25 base pairs from exon boundaries). For every single position (nucleotide), median conservation scores were calculated by deeptools [2] over a set of non-redundant exons (different in their genomic coordinates) present in the collapsed projected Iso-seq models (hg38) (top). PhastCons conservation scores for every single position across these exons are represented as a color gradient (bottom) according to the color legend (right). Only exons that are 100-500 nucleotides long were kept for this representation (N=84,785 exons). Conservation scores were retrieved from 17-way alignments (UCSC).

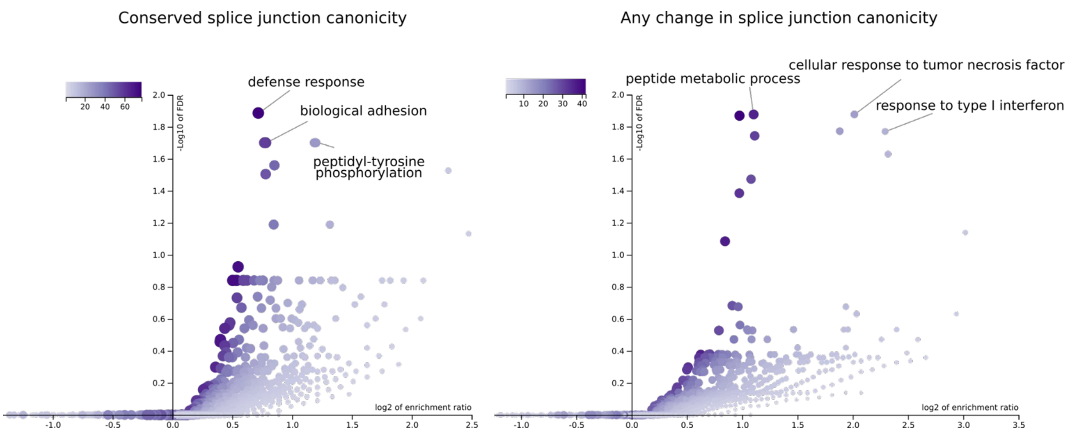


**Supplementary Figure 13.** Functional enrichment for genes expressing species-specific transcripts that arise from conserved splice site canonicity (left, N=627 genes) and not conserved splice site canonicity (right, N=280 genes) across primates. The enrichment was computed by over-representation analysis (ORA) using the Gene Ontology Biological Process database (affinity propagation clustering). False discovery rates (FDR) were adjusted using the Benjamini-Hochberg method [3]. Color legend indicates the number of overlapping genes between the gene set under evaluation and the pathway gene set.

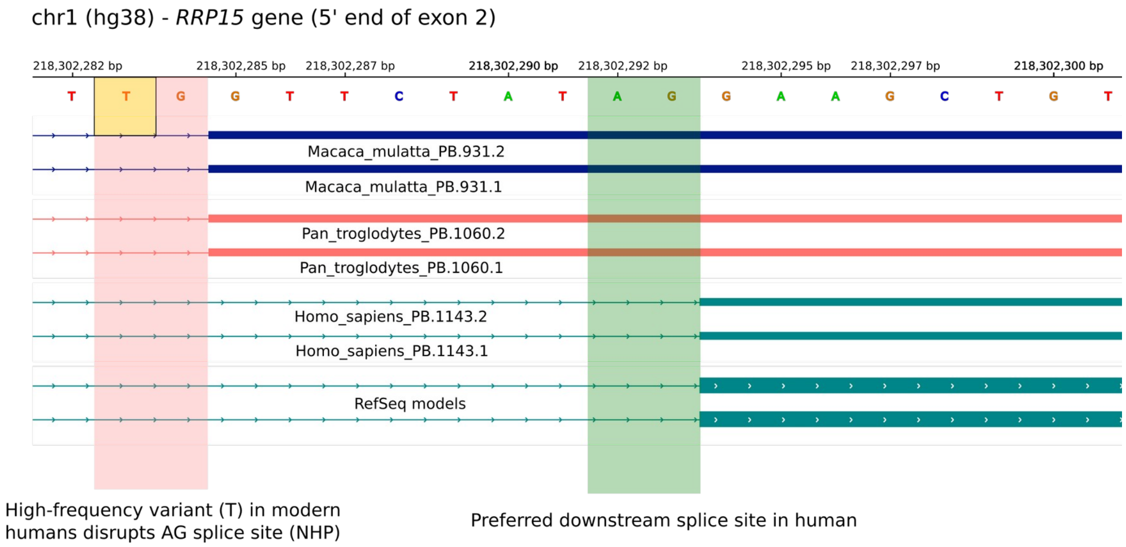


**Supplementary Figure 14.** Example of a gene**,** *RRP15*, with a human-specific splice site mutation altering the definition of the 5’ boundary of *RRP15* exon 2. *Macaca mulatta* (blue) and *Pan troglodytes* (pink) Iso-seq projected isoforms are shown together with *Homo sapiens* Iso-seq isoforms and RefSeq models in hg38 assembly (green).

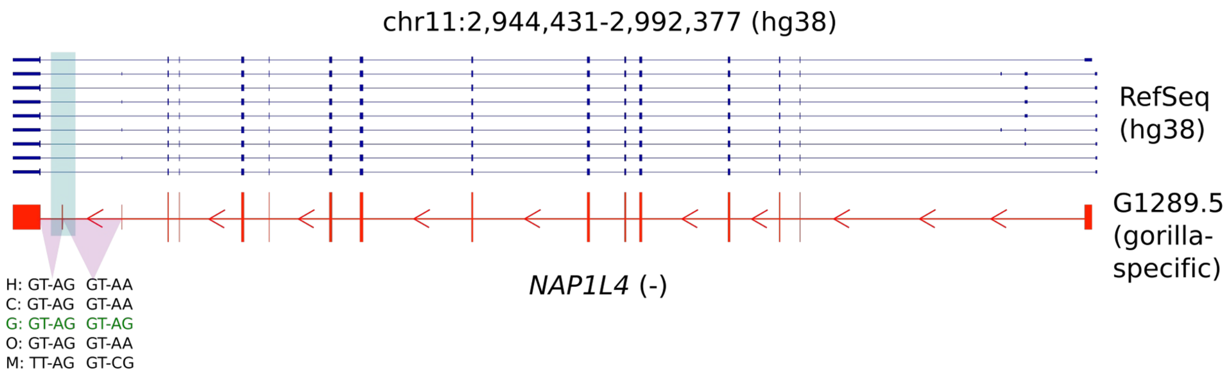


**Supplementary Figure 15.** Example of a gene, *NAP1L4*, with a gorilla-specific splice site mutation resulting in an exonization event. RefSeq has already annotated this gorilla-specific 3' UTR exon in gorGor6 assembly.

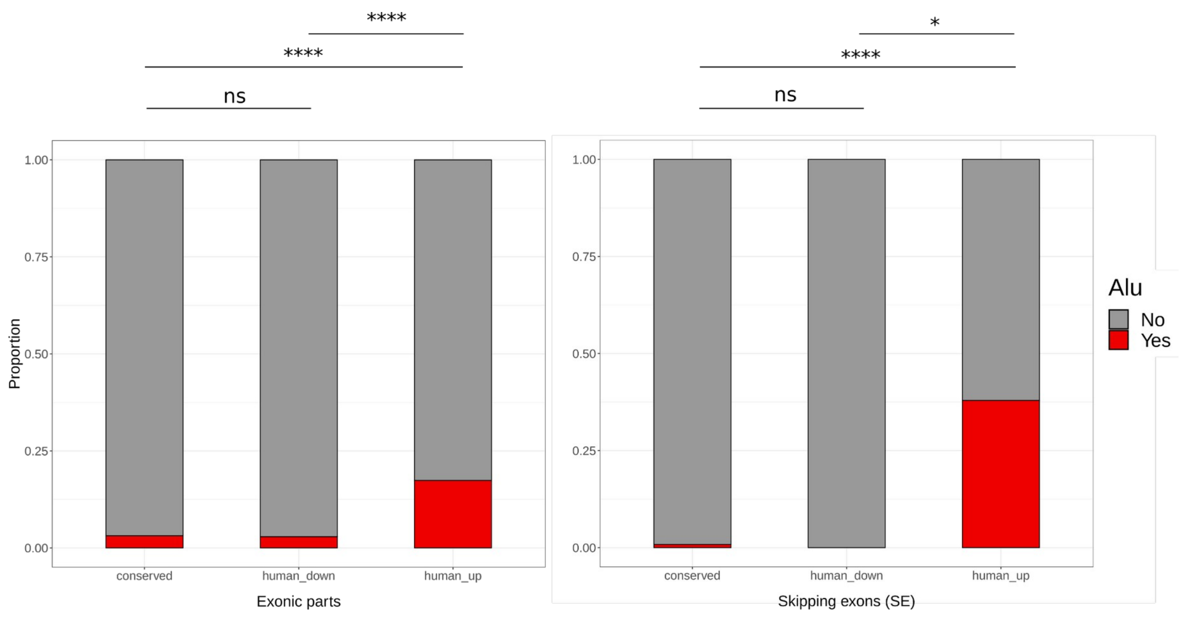


**Supplementary Figure 16.** Proportion of exonic parts intersecting Alu elements (conserved usage: 'conserved'; human-specific downregulated: 'human_down'; human-specific upregulated: 'human_up'). All exonic parts are shown to the left while skipping exons (SE) are displayed to the right. Alu elements were retrieved from hg38 repeatMasker track (UCSC). Statistical significance in the difference in proportions across groups was evaluated using Fisher's exact test. P-values were adjusted using Bonferroni's method (**** = P ≤ 0.0001, * = P ≤ 0.05, ns = P > 0.05).

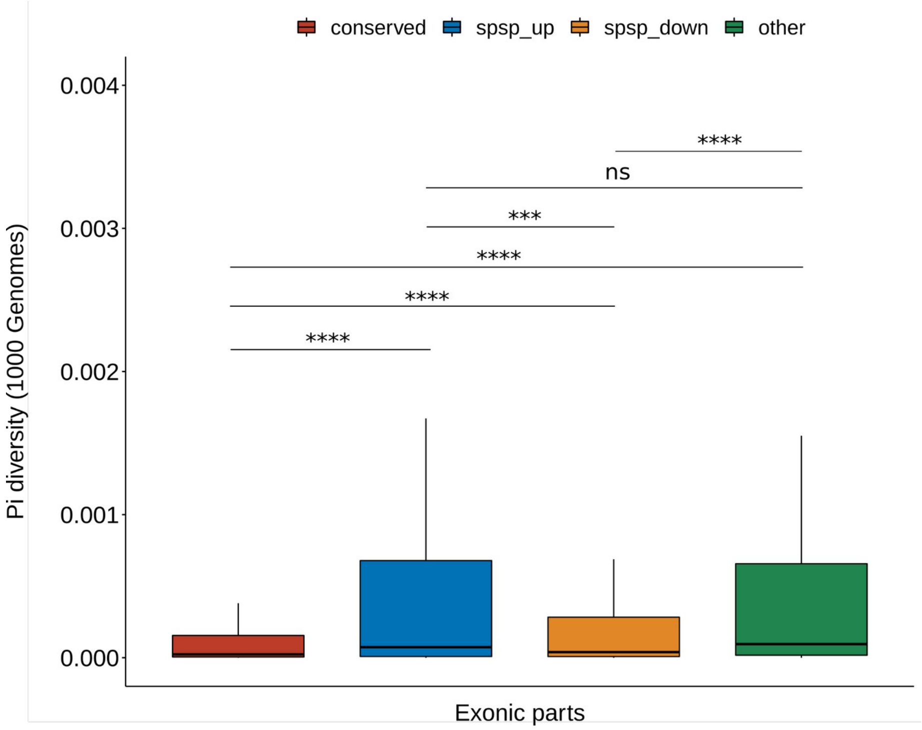


**Supplementary Figure 17.** Average nucleotide diversity (pi) in human populations for exonic parts with conserved usage (‘conserved’, N=95,453), exonic parts with species-specific upregulation (‘spsp_up’, N=941) and downregulation (‘spsp_down’, N=1,340), and exonic parts showing other usage changes (‘other’, N=277). ‘other’ usage changes correspond to usage differences between groups of species (*e.g.,* any 2 species show differential usage *versus* the remaining 3 species). Pi diversity estimates were calculated from the 1000 Genomes data mapped against hg38. Only exonic parts longer than 5 bases with all nucleotides included as passed bases according to 1000 Genomes strict mask are shown. Statistical significance of the difference across groups was assessed by Wilcoxon test and adjusted by Holm method (**** = P ≤ 0.0001, *** = P ≤ 0.001, ns = P > 0.05).

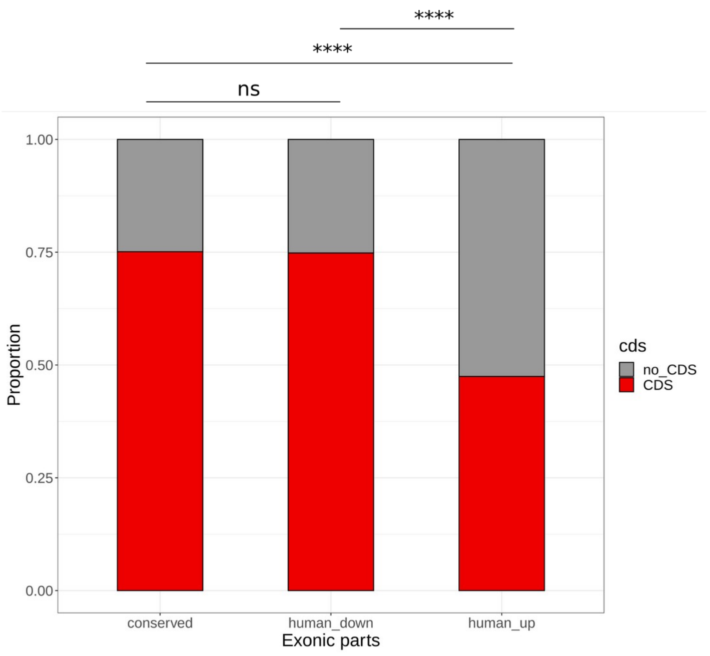


**Supplementary Figure 18.** Proportion of exonic parts intersecting coding regions (CDS) predicted from Iso-seq projected models (conserved usage: ‘conserved’; human-specific downregulated: ‘human_down’; human-specific upregulated: ‘human_up’). Statistical significance in the difference in proportions across groups was evaluated using Fisher’s exact test. P-values were adjusted using Bonferroni’s method (**** = P ≤ 0.0001, ns = P > 0.05).

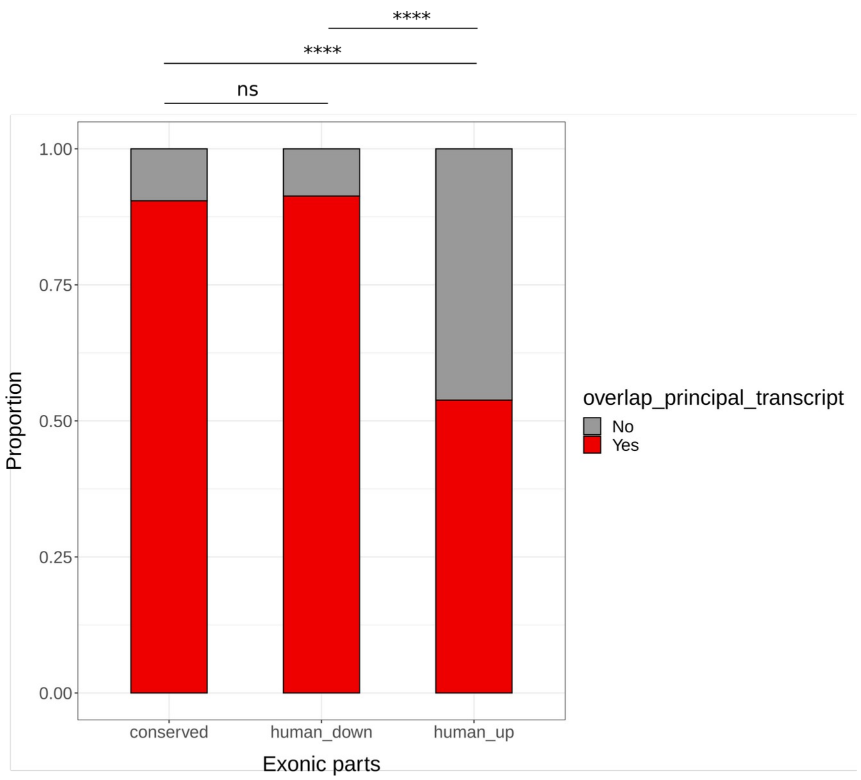


**Supplementary Figure 19.** Proportion of exonic parts intersecting regions included in APPRIS principal transcripts (hg38) (conserved usage: ‘conserved’; human-specific downregulated: ‘human_down’; human-specific upregulated: ‘human_up’). Statistical significance in the difference in proportions across groups was evaluated using Fisher’s exact test. P-values were adjusted using Bonferroni’s method (**** = P ≤ 0.0001, ns = P > 0.05).

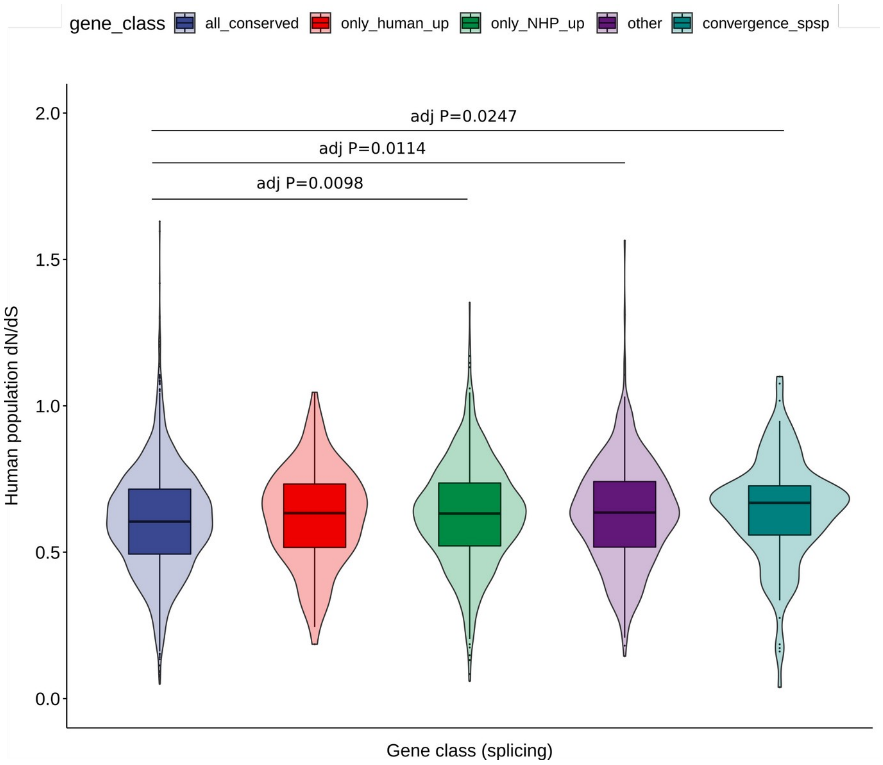


**Supplementary Figure 20.** Ratio of nonsynonymous to synonymous substitutions (dN/dS) in human populations for each of the gene classes according to their splicing and usage patterns. Statistical significance for the pairwise comparisons was obtained using the Dwass-Steel-Critchlow-Fligner all-pairs test. P-values were adjusted using the single-step method. Only adjusted p-values lower than 0.05 are shown.

### Supplementary Tables

**Supplementary Table 1.** Statistics of PacBio Iso-seq and Illumina RNA-seq data production in 5 primate species. CCS: circular consensus sequence; FLNC: full-length non-chimeric reads.

| Data production | | Human | Chimpanzee | Gorilla | Orangutan | Macaque |
|---|---|---|---|---|---|---|
| Iso-seq | CCS | 2,901,238 | 2,478,756 | 2,144,402 | 1,959,837 | 2,226,668 |
| | FLNC | 2,137,334 | 1,832,946 | 1,564,808 | 1,213,522 | 1,458,642 |
| | Polished transcript | 104,486 | 98,602 | 86,768 | 64,693 | 83,262 |
| | Non-redundant transcript | 63,577 | 55,417 | 46,191 | 34,124 | 48,247 |
| | High-quality transcript | 50,769 | 34,871 | 26,003 | 13,766 | 23,201 |
| RNA-seq | Read | 331,829,884 | 472,536,800 | 499,997,165 | 430,203,432 | 495,996,165 |
| | Junction | 488,229 | 607,996 | 546,444 | 396,136 | 488,229 |

**Supplementary Table 2.** Number of Iso-seq isoforms in each SQANTI structural category according to Ensembl models (V91). Only isoforms passing SQANTI QC (not artifacts) are shown. FSM: full splice match; ISM: incomplete splice match; NIC: novel in catalog; NNC: novel not in catalog. A detailed description of SQANTI structural categories can be found in [1].

| Species | FSM | ISM | NIC | NNC | antisense | fusion | genic | genic intron | intergenic |
|---|---|---|---|---|---|---|---|---|---|
| Human | 25,165 | 3,625 | 16,647 | 3,831 | 184 | 51 | 465 | 543 | 258 |
| Chimpanzee | 13,267 | 2,454 | 9,735 | 7,055 | 244 | 24 | 726 | 480 | 886 |
| Gorilla | 10,889 | 1,656 | 6,343 | 5,245 | 190 | 17 | 530 | 293 | 840 |
| Orangutan | 6,371 | 1,010 | 2,682 | 2,515 | 110 | 11 | 294 | 167 | 606 |
| Macaque | 9,257 | 1,755 | 6,056 | 4,047 | 168 | 23 | 535 | 412 | 948 |

**Supplementary Table 3.** Number of alternative splicing events detected in Iso-seq transcriptomes. SE: skipping exons; RI: retained introns; A5SS: alternative 5’ splice sites; A3SS: alternative 3’ splice sites; MX: mutually exclusive exons.

| Species | Total | SE | RI | A5SS | A3SS | MX |
|---|---|---|---|---|---|---|
| Human | 16,337 | 6,401 | 4,062 | 2,845 | 2,658 | 371 |
| Chimpanzee | 10,371 | 4,233 | 2,529 | 1,650 | 1,716 | 243 |
| Gorilla | 7,312 | 2,892 | 1,728 | 1,307 | 1,241 | 144 |
| Orangutan | 3,015 | 1,280 | 622 | 586 | 462 | 65 |
| Macaque | 6,144 | 2,466 | 1,506 | 1,048 | 994 | 130 |

**Supplementary Table 4.** Number of detected peptides per species according to their presence in the simulated digestion of Iso-seq isoforms projected to each genome (plausible in genome), UniProt and RefSeq reference proteomes. Detected peptides must satisfy FDR<5%, Mascot IonScore > 20, reporter ion intensity signal (abundance) per sample > 50 (in any sample of a given species), and species-wise median ratio of sample abundance to pool abundance > 0.6 (excluding matches to common contaminant sequences and peptides arising from more than 1 tryptic miscleavage). The numbers correspond to distinct tryptic peptides considering isoleucine and leucine (I/L) as indistinguishable by mass spectrometry.

| Species | Detected peptides | Detected peptides and plausible in genome | Novel peptides in UniProt and plausible in genome | Novel peptides in RefSeq and plausible in genome |
|---|---|---|---|---|
| Human | 24,308 | 24,180 | 22 | 25 |
| Chimpanzee | 25,158 | 24,733 | 320 | 98 |
| Gorilla | 25,577 | 24,409 | 882 | 94 |
| Orangutan | 25,741 | 25,205 | 1,590 | 225 |
| Macaque | 23,743 | 23,299 | 207 | 75 |

**Supplementary Table 5.** Functional enrichment for genes displaying species-specific transcript gains (over-representation analysis using Gene Ontology Biological Process database in WebGestalt, FDR ≤ 0.05).

| Gene Set | Description | Size | Expect | Enrichment ratio | P-value | FDR |
|---|---|---|---|---|---|---|
| GO:0006952 | defense response | 434 | 64.315 | 1.6481 | 2.32E-08 | 0.00008551 |
| GO:0045087 | innate immune response | 270 | 40.012 | 1.7495 | 6.90E-07 | 0.0010283 |
| GO:0031347 | regulation of defense response | 231 | 34.232 | 1.8112 | 8.68E-07 | 0.0010283 |
| GO:0018108 | peptidyl-tyrosine phosphorylation | 109 | 16.153 | 2.2287 | 1.1184E-06 | 0.0010283 |
| GO:0018212 | peptidyl-tyrosine modification | 110 | 16.301 | 2.2084 | 1.4321E-06 | 0.0010535 |
| GO:0034341 | response to interferon-gamma | 67 | 9.9288 | 2.4172 | 0.000015048 | 0.0092242 |
| GO:0002252 | immune effector process | 456 | 67.575 | 1.465 | 0.000022508 | 0.011826 |
| GO:0032101 | regulation of response to external stimulus | 203 | 30.083 | 1.7286 | 0.00002846 | 0.013085 |
| GO:0031349 | positive regulation of defense response | 146 | 21.636 | 1.8488 | 0.000048233 | 0.016607 |
| GO:0050776 | regulation of immune response | 325 | 48.162 | 1.5365 | 0.000052747 | 0.016607 |
| GO:0045088 | regulation of innate immune response | 142 | 21.043 | 1.8533 | 0.000056417 | 0.016607 |
| GO:0071346 | cellular response to interferon-gamma | 59 | 8.7433 | 2.4018 | 0.000057105 | 0.016607 |
| GO:0002684 | positive regulation of immune system process | 337 | 49.94 | 1.5218 | 0.000058697 | 0.016607 |
| GO:0050778 | positive regulation of immune response | 254 | 37.641 | 1.594 | 0.000093189 | 0.024482 |
| GO:0050707 | regulation of cytokine secretion | 55 | 8.1505 | 2.3311 | 0.00020184 | 0.046466 |
| GO:0019221 | cytokine-mediated signaling pathway | 235 | 34.825 | 1.5793 | 0.00023632 | 0.046466 |
| GO:0002682 | regulation of immune system process | 477 | 70.687 | 1.3864 | 0.0002377 | 0.046466 |
| GO:0034340 | response to type I interferon | 43 | 6.3722 | 2.5109 | 0.00024081 | 0.046466 |
| GO:0060333 | interferon-gamma-mediated signaling pathway | 39 | 5.7795 | 2.5954 | 0.00024577 | 0.046466 |

| | | | | | | |
|---|---|---|---|---|---|---|
| GO:0001816 | cytokine production | 252 | 37.344 | 1.5531 | 0.00025768 | 0.046466 |
| GO:0001817 | regulation of cytokine production | 226 | 33.491 | 1.5825 | 0.00029057 | 0.046466 |
| GO:0022610 | biological adhesion | 313 | 46.384 | 1.4876 | 0.00026845 | 0.046466 |
| GO:0071345 | cellular response to cytokine stimulus | 353 | 52.312 | 1.4528 | 0.00028704 | 0.046466 |

**Supplementary Table 6.** Functional enrichment for genes expressing transcripts which show a change in junction canonicity in any species (over-representation analysis using Panther pathways database in WebGestalt, FDR ≤ 0.05).

| Gene Set | Description | Size | Expect | Enrichment ratio | P-value | FDR |
|---|---|---|---|---|---|---|
| P00010 | B cell activation | 47 | 6.0893 | 2.4633 | 0.00038484 | 0.01917 |
| P00053 | T cell activation | 44 | 5.7006 | 2.4559 | 0.00063901 | 0.01917 |
| P00038 | JAK/STAT signaling pathway | 11 | 1.4252 | 4.2101 | 0.0011021 | 0.022041 |

**Supplementary Table 7.** Functional enrichment for genes displaying species-specific exon usage changes (over-representation analysis using Gene Ontology Biological Process database in WebGestalt, FDR ≤ 0.05).

| Gene Set | Description | Size | Expect | Enrichment Ratio | P-value | FDR |
|---|---|---|---|---|---|---|
| GO:0033365 | protein localization to organelle | 486 | 99.542 | 1.5471 | 1.07E-09 | 4.6988E-06 |
| GO:0022613 | ribonucleoprotein complex biogenesis | 268 | 54.892 | 1.6942 | 2.37E-08 | 0.000045672 |
| GO:0034660 | ncRNA metabolic process | 330 | 67.59 | 1.6127 | 3.11E-08 | 0.000045672 |
| GO:0042254 | ribosome biogenesis | 162 | 33.181 | 1.8686 | 1.08E-07 | 0.00011924 |
| GO:0006281 | DNA repair | 329 | 67.386 | 1.573 | 2.03E-07 | 0.0001792 |
| GO:0140053 | mitochondrial gene expression | 101 | 20.687 | 2.0303 | 9.71E-07 | 0.00071234 |
| GO:0032543 | mitochondrial translation | 85 | 17.41 | 2.1253 | 1.1317E-06 | 0.00071234 |
| GO:0034470 | ncRNA processing | 224 | 45.88 | 1.6347 | 2.6492E-06 | 0.00111 |
| GO:0060337 | type I interferon signaling pathway | 44 | 9.012 | 2.5521 | 2.7976E-06 | 0.00111 |
| GO:0071357 | cellular response to type I interferon | 44 | 9.012 | 2.5521 | 2.7976E-06 | 0.00111 |
| GO:0016072 | rRNA metabolic process | 143 | 29.289 | 1.8095 | 0.000002852 | 0.00111 |

| | | | | | | |
|---|---|---|---|---|---|---|
| GO:0006364 | rRNA processing | 122 | 24.988 | 1.8809 | 3.0231E-06 | 0.00111 |
| GO:0006403 | RNA localization | 134 | 27.446 | 1.8218 | 4.3445E-06 | 0.0014724 |
| GO:0016032 | viral process | 406 | 83.157 | 1.4431 | 4.8853E-06 | 0.0015375 |
| GO:0045087 | innate immune response | 352 | 72.096 | 1.4703 | 7.2226E-06 | 0.0019632 |
| GO:0034340 | response to type I interferon | 49 | 10.036 | 2.3914 | 0.000007493 | 0.0019632 |
| GO:0072594 | establishment of protein localization to organelle | 265 | 54.277 | 1.5476 | 7.9216E-06 | 0.0019632 |
| GO:0007005 | mitochondrion organization | 269 | 55.096 | 1.5428 | 8.0204E-06 | 0.0019632 |
| GO:0043604 | amide biosynthetic process | 405 | 82.952 | 1.4225 | 0.000012415 | 0.0028791 |
| GO:0044403 | symbiont process | 431 | 88.277 | 1.4047 | 0.000014448 | 0.0031829 |
| GO:0043043 | peptide biosynthetic process | 339 | 69.434 | 1.4114 | 0.000095917 | 0.017321 |
| GO:0022618 | ribonucleoprotein complex assembly | 142 | 29.084 | 1.6504 | 0.00012726 | 0.019658 |
| GO:0050776 | regulation of immune response | 442 | 90.53 | 1.3476 | 0.00012939 | 0.019658 |
| GO:0070925 | organelle assembly | 433 | 88.687 | 1.3418 | 0.00019047 | 0.025431 |
| GO:0050658 | RNA transport | 115 | 23.554 | 1.6982 | 0.00023056 | 0.029024 |
| GO:0006397 | mRNA processing | 307 | 62.88 | 1.3995 | 0.00029714 | 0.035384 |
| GO:0055086 | nucleobase-containing small molecule metabolic process | 362 | 74.145 | 1.3622 | 0.00032404 | 0.037571 |
| GO:0016071 | mRNA metabolic process | 432 | 88.482 | 1.3223 | 0.00040547 | 0.043573 |
| GO:0009117 | nucleotide metabolic process | 316 | 64.723 | 1.3751 | 0.0005213 | 0.049971 |
| GO:0000723 | telomere maintenance | 93 | 19.048 | 1.7324 | 0.00052696 | 0.049971 |
| GO:0032200 | telomere organization | 93 | 19.048 | 1.7324 | 0.00052696 | 0.049971 |
| GO:0051640 | organelle localization | 354 | 72.506 | 1.3516 | 0.00053319 | 0.049971 |
| GO:0051642 | centrosome localization | 17 | 3.4819 | 2.872 | 0.00059298 | 0.049971 |

**Supplementary Table 8.** Tandem mass tag (TMT) labeling reagents used in each of the samples.

| Sample name | TMT channel | Species | Code |
|---|---|---|---|
| Mix-1 | 126 | Human | GM19238 |
| Mix-1 | 127 | Chimpanzee | CH507 |
| Mix-1 | 128 | Gorilla | DIAN |
| Mix-1 | 129 | Macaque | R02027 |
| Mix-1 | 130 | Orangutan | PPY6 |
| Mix-1 | 131 | Pool | - |
| Mix-2 | 126 | Pool | - |
| MIx-2 | 127 | Human | GM12878 |
| Mix-2 | 128 | Chimpanzee | CH170 |
| Mix-2 | 129 | Gorilla | OMOYE |
| Mix-2 | 130 | Macaque | R05040 |
| Mix-2 | 131 | Orangutan | EB185(JC) |
| Mix-3 | 126 | Orangutan | CRL-1850 (PUTI) |
| Mix-3 | 127 | Pool | - |
| Mix-3 | 128 | Human | GM19150 |
| Mix-3 | 129 | Chimpanzee | CH322 |
| Mix-3 | 130 | Gorilla | GG05 |
| Mix-3 | 131 | Macaque | R94011 |

### Supplementary Data

**Supplementary Data 1.** SQANTI classification files for Iso-seq transcripts passing SQANTI quality filtering. Genomic coordinates are based on hg38, panTro5, gorGor4, ponAbe2 and rheMac8.

**Supplementary Data 2.** All detected peptides by mass spectrometry experiments.

**Supplementary Data 3.** Detected novel peptides according to RefSeq annotations. Isoleucine and leucine amino acids are designed as ‘Z’ since they are indistinguishable by mass spectrometry experiments.

**Supplementary Data 4.** GFF file for the projected isoform models in hg38 coordinates.

**Supplementary Data 5.** SQANTI classification file for the projected isoform models in hg38.

**Supplementary Data 6.** Species-specific exon gains. Genomic coordinates are based on hg38, panTro6, gorGor6, ponAbe3 and rheMac10.

**Supplementary Data 7.** Expression matrix for projected isoform models across samples (TPM, including batch effect correction and TMM normalization).

**Supplementary Data 8.** Binary expression matrix (1=presence, 0=absence) for projected isoform models across species. Only transcripts showing consistent expression in all samples from the same species are included.

**Supplementary Data 9.** Classification of projected isoform models according to their isoform usage patterns.

**Supplementary Data 10.** Classification of exonic parts according to differential exon usage (DEU) results in hg38 coordinates.

### Supplementary References


1. Tardaguila, M. *et al.* SQANTI: extensive characterization of long-read transcript sequences for quality control in full-length transcriptome identification and quantification. *Genome Res.* **28**, 396–411 (2018).
2. Ramírez, F. *et al.* deepTools2: a next generation web server for deep-sequencing data analysis. *Nucleic Acids Res.* **44**, W160–W165 (2016).
3. Benjamini, Y., Drai, D., Elmer, G., Kafkafi, N. & Golani, I. Controlling the false discovery rate in behavior genetics research. *Behav. Brain Res.* **125**, 279–284 (2001).
